# Epididymis-specific RNase A family genes regulate fertility and small RNA processing

**DOI:** 10.1101/2024.08.26.608813

**Authors:** Joshua F. Shaffer, Alka Gupta, Geetika Kharkwal, Edgardo E. Linares, Andrew D. Holmes, Sol Katzman, Upasna Sharma

## Abstract

Sperm small RNAs are implicated in intergenerational transmission of paternal environmental effects. Small RNAs generated by cleavage of tRNAs, known as tRNA fragments (tRFs), are an abundant class of RNAs in mature sperm, and can be modulated by environmental conditions. The ribonuclease(s) responsible for the biogenesis of tRFs in the male reproductive tract remains unknown. Angiogenin, a member of the Ribonuclease A superfamily (RNase A), cleaves tRNAs to generate tRFs in response to cellular stress. Four paralogs of Angiogenin, namely *Rnase9*, *Rnase10, Rnase11*, and *Rnase12*, are specifically expressed in the epididymis—a long, convoluted tubule where sperm mature and acquire fertility and motility. The biological functions of these genes remain largely unknown. Here, by generating mice deleted for all four genes (*Rnase9-12-/-*, termed “KO” for Knock Out), we report that these genes regulate fertility and RNA processing. KO mice showed complete male sterility. KO sperm fertilized oocytes *in vitro* but failed to efficiently fertilize oocytes *in vivo,* likely due to an inability of sperm to pass through the utero-tubular junction. Intriguingly, there were decreased levels of fragments of tRNAs (tRFs) and rRNAs (rRNA-derived small RNAs or rsRNAs) in the KO epididymis and epididymal luminal fluid, implying that *Rnase9-12* regulate the biogenesis and/or stability of tRFs and rsRNAs. Importantly, KO sperm showed a dramatic decrease in the levels of tRFs, demonstrating a role of *Rnase9-12* in regulating sperm RNA composition. Together, our results reveal an unexpected role of four epididymis-specific non-canonical RNase A family genes in fertility and RNA processing.

## INTRODUCTION

Sperm released from the testicular seminiferous tubules are not capable of fertilization; they acquire forward motility and fertilization capacity during post-testicular maturation in the epididymis [1]. The epididymis is anatomically and functionally subdivided into three broad segments: caput (proximal), corpus (middle), and cauda (distal) epididymis. Mature sperm are stored in the cauda epididymis and released upon copulation via the vas deferens. Sperm encounter a variety of proteins, lipids, and ions in the epididymal luminal microenvironment, which is shaped by the secretory activity of the local epithelial cells [2–5]. While it is well established that the epididymis plays an essential role in post-testicular sperm maturation and fertility [1, 6, 7], the specific factors in the epididymal microenvironment that regulate fertility are poorly defined.

The epididymis also plays a role in shaping the RNA composition of mature sperm. The RNA payload of mature sperm is composed of a diverse class of small RNAs, including microRNAs (miRNAs), piwi-interacting RNAs (piRNAs), and cleavage products of transfer RNAs (tRNA fragments or tRFs) and ribosomal RNAs (rRNA-derived small RNAs or rsRNAs). Recent studies revealed that sperm undergo dramatic changes in their small RNA payload during epididymal transit; while testicular sperm are highly enriched in piRNAs, mature sperm in the cauda epididymis are enriched in tRFs and rsRNAs [8–12]. We and others reported that a subset of these newly acquired small RNAs are delivered to sperm from epididymal epithelial cells [8, 9, 12–17]. How tRFs and rsRNAs are generated in the epididymal epithelial cells remains unknown. Importantly, sperm small RNAs have been implicated in intergenerational transmission of paternal environmental effects [18–22]. Small RNAs in mature sperm are altered in response to environmental conditions and delivered to the zygote at fertilization, where they can modulate early embryonic development. Hence, it is crucial to elucidate the biogenesis of sperm small RNAs, as it may be a target of signaling pathways that link paternal environment conditions to offspring phenotypes.

tRNA cleavage has been characterized in several biological contexts, where it is typically induced in response to stress conditions. In budding yeast, *Tetrahymena thermophilia*, and *Arabidopsis*, RNase T2 family endonucleases process tRNAs [23–25], whereas in mammalian cells exposed to stress, the RNase A family member Angiogenin (encoded by *Rnase5*) cleaves tRNAs [26–28]. RNase A and T2 family endonucleases leave a 2’-3’ cyclic phosphate (2’3’cP) or a 3’ phosphate (3’P) at the 3’ end of RNA, both of which can interfere with small RNA cloning protocols. T4 polynucleotide kinase (PNK) removes cyclic phosphates from the 3’ ends of RNA molecules when the reaction is carried out in the absence of ATP. We previously reported that treatment of sperm total RNA with PNK allows adapter ligation and cloning for deep sequencing of RNAs with a 2’3’cP or 3’P at the 3’ end, revealing an abundant class of 5’ tRFs of 33-36 nt.

These tRFs are slightly longer than the 28-32 nt tRFs sequenced without PNK treatment. The population of longer tRFs was typically generated by cleavage within or immediately downstream of the anticodon. These data suggested that the longer tRFs are likely the initial cleavage products generated by an RNase A or RNase T2 family member at the accessible tRNA loop structures, with the previously described shorter (28-32 nt) fragments representing secondary degradation or trimmed products [8]. In addition, we found that PNK treatment resulted in a dramatic increase in the levels of rsRNAs in sperm and that rsRNAs were the most abundant class of small RNAs in sperm [8]. Overall, our previous studies revealed that tRFs and rsRNAs are generated by RNase A and /or RNase T2 endonuclease in the male reproductive tract. Indeed, recent studies showed that inflammation and stress can induce Angiogenin [29] and RNase T2 [30] expression in the epididymis and increase levels of specific tRFs in sperm. However, robust tRNA cleavage is observed in the epididymis even in the absence of stress [9]. The endonuclease(s) responsible for the biogenesis of tRFs and rsRNAs in the male reproductive tract under non-stress, physiological conditions remain unknown.

The RNase A superfamily is a vertebrate-specific gene family and consists of 13 members. While proteins within this superfamily share structural and catalytic properties, their primary sequences have undergone considerable divergence, presumably to facilitate the emergence of novel functions [31]. RNases 1-8 are known as canonical members due to the presence of conserved catalytic residues and disulfide bonds, and these RNases exhibit diverse expression patterns and catalytic activity [32]. Functionally, RNases 1-8 regulate catalytic activity-dependent and independent processes, including RNA degradation, antimicrobial and antiviral activity, and innate immunity [27, 33–36]. In addition to RNases 1-8, another four RNase A paralogs — RNases 9-12 —, are highly and specifically expressed in different segments of the epididymis [32, 37–42]. Genetic studies of *Rnase10* and *Rnase9* indicate a functional role of these genes in male reproduction. Deletion of *Rnase10* resulted in a male fertility defect in the C57BL/6 strain background but not in CD1 mice, thus showing a strain-specific phenotype [43]. *Rnase9* knockout mice were fertile but showed slightly reduced sperm motility immediately after release from the epididymis [44]. Additionally, human RNase 9 localizes to sperm heads, and recombinant RNase 9 has antimicrobial activity [42]. Whether RNase 11 and RNase 12 play a role in male fertility has not been investigated.

RNases 9-12 share 15-30% sequence identity with the canonical RNases and retain the signal peptide and three conserved disulfide bonds. Due to a lack of conserved active site sequence motifs, they are predicted to be catalytically inactive and are known as non-canonical members of the RNase A family [32]. A bacterial RNase was recently discovered that adopts the same structural folds as Angiogenin and its paralogs but has no sequence similarity with RNase A family members and lacks the conserved disulfide bonds and catalytic triad characteristic of RNase A family [45]. This protein was shown to possess potent RNase activity *in vitro* and *in vivo* despite having a catalytic core distinct from that found in the canonical RNase A enzymes. Moreover, Angiogenin has very weak catalytic activity (10^5^-10^6^ times lower than that of RNase 1) but can still cleave tRNAs [46, 47]. These observations suggest that RNases 9-12, which are paralogs of Angiogenin, could have catalytic activity and warrant examination of the role of these proteins in RNA processing in the male reproductive tract.

Here, we deleted the genomic cluster on chromosome 14 harboring the *Rnase9, Rnase10, Rnase11, and Rnase12* genes in the FVB mouse strain background. The *Rnase9-12* knockout (KO) male mice showed complete loss of fertility as they failed to generate pups when mated with wild-type females. While KO sperm were capable of fertilizing oocytes *in vitro* and did not show any defects in morphology and number, KO sperm had undetectable levels of the peptidase protein ADAM3. Loss of ADAM3 from mature sperm has been shown to cause a defect in traversing through the utero-tubular junction, suggesting that KO sperm likely cannot reach the oocytes to fertilize. We found that the epididymis of KO mice had transcriptomic and proteomic changes consistent with fertility defects. Next, we investigated whether these proteins are involved in RNA processing in the epididymis. Indeed, tRFs and rsRNAs were downregulated in the epididymis and sperm of KO mice, demonstrating that *Rnase9-12* play a role in regulating small RNA levels in the epididymis and, thus, modulate the repertoire of small RNAs in mature sperm.

## RESULTS

### Deletion of *the Rnase9-12* gene cluster resulted in loss of male fertility

To study the role of *Rnase9-12* genes in the epididymis, we generated a mouse line with deletion of the genomic cluster harboring these four genes using CRISPR/Cas9 genome editing **(Figure S1A).** The heterozygous (*Rnase9-12^-/+^* or HET) and homozygous (*Rnase9-12^-/-^* or KO) deletion mice showed the expected decrease in the levels of transcripts of these genes in the epididymis when compared to wild-type (*Rnase9-12^+/+^* or WT), as seen by mRNA sequencing and quantitative real time-PCR (**Figure 1A-D and Figure S1B).** These data confirmed a complete knockout of the *Rnase9-12* locus. Moreover, our mRNA-seq analysis corroborated previous studies reporting segment-specific expression of *Rnase9-12*, with *Rnase10* being expressed exclusively in the caput epididymis [38, 40, 43]. We monitored the genotype of mice in all subsequent experiments to ensure the knockout of all four genes.

**Figure 1:**
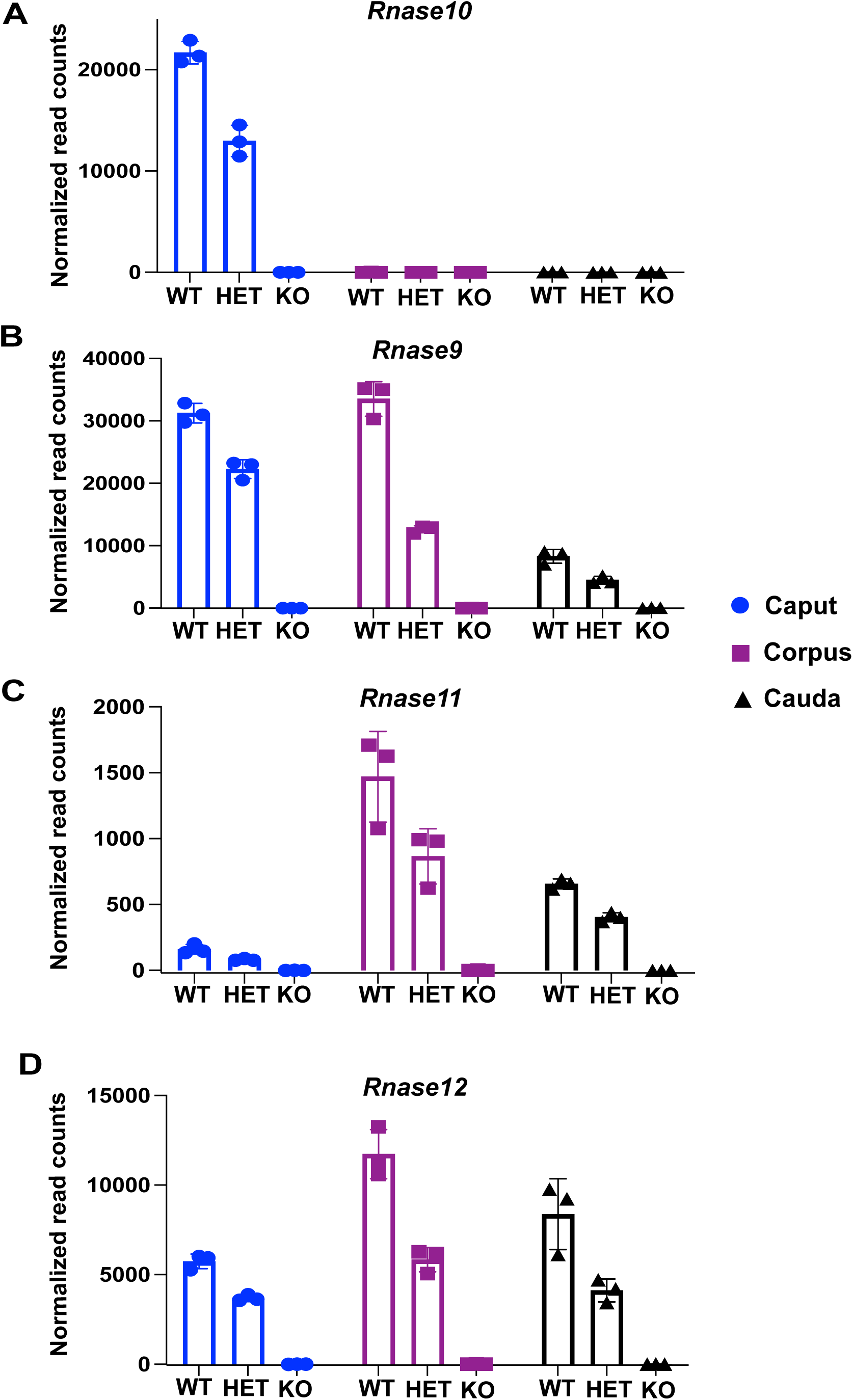
Generation of mice with deletion of a genomic locus harboring four reproductive-tract-specific RNase A family genes. Normalized read counts of *Rnase10* (A), *Rnase9* (B), *Rnase11* (C), and *Rnase12* (D) from mRNA-seq performed on the caput, corpus, and cauda epididymis tissues of wild-type (WT), heterozygous deletion (HET) and homozygous deletion (KO) males. Normalized read counts were obtained using DESeq2 analysis, and bar graphs represent read counts from three biological replicates.

At the phenotypic level, the KO mice do not show any differences in physical appearance, behavior, or weight compared to WT mice. Histology of reproductive tract tissues showed no differences in the overall morphology in KO mice compared to WT mice (data not shown). To investigate potential functional roles for these genes in reproduction, we next mated WT, HET, or KO males with WT females and vice versa. In matings with WT females, WT and HET males produced normal numbers of pups, while KO males failed to produce any pups; in matings with WT males, females of all three genotypes produced normal numbers of pups (**Figure 2A-C**). Over 4-6 months, the KO males did not produce any litters (n=8 males). In contrast, HET males showed normal reproductive capacity, demonstrating the haplosufficiency of these four genes. These data indicate that the *Rnase9-12* genomic cluster is required for male fertility.

**Figure 2:**
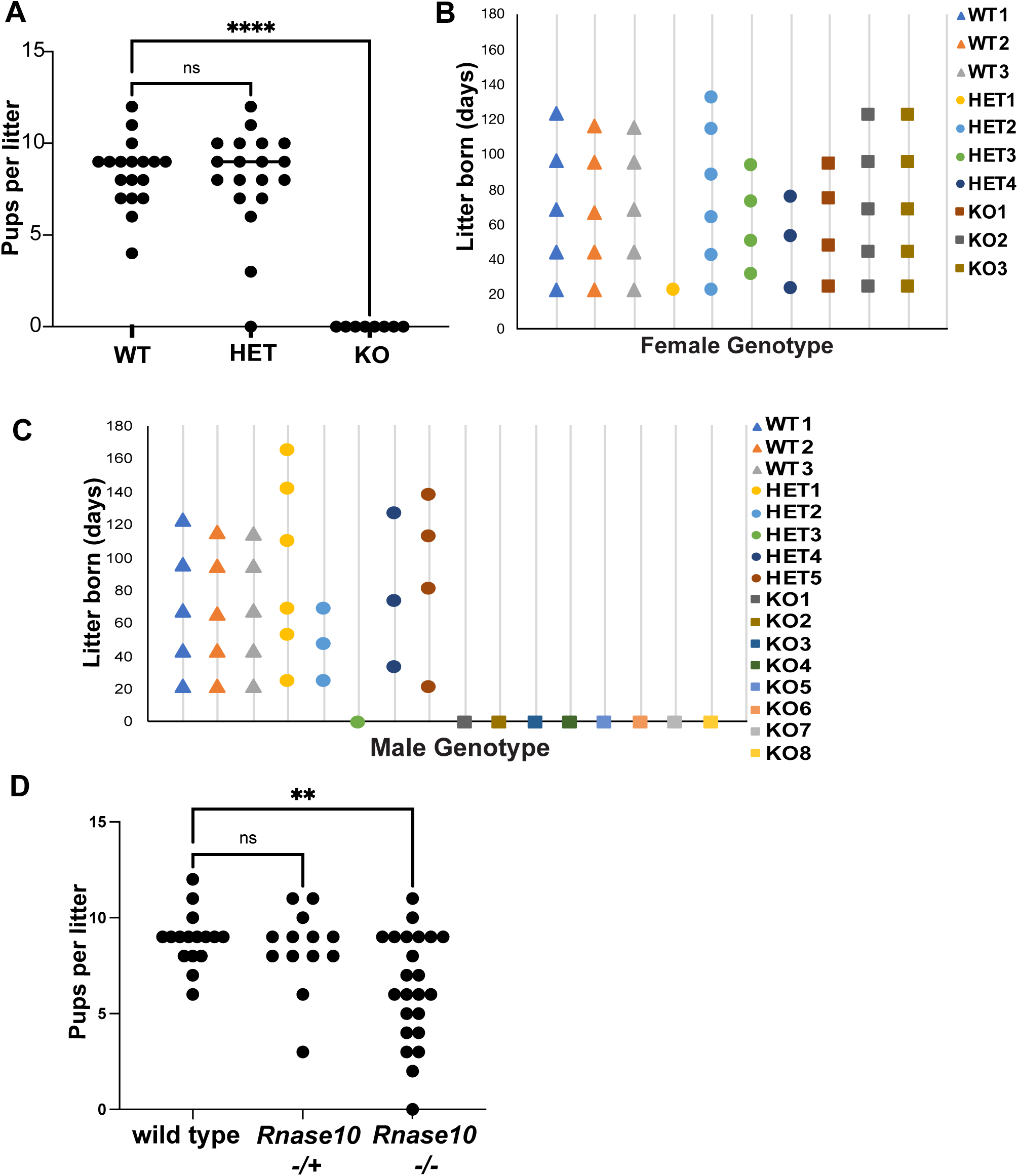
Loss of male fertility in *Rnase9-12* KO mice. A) Number of pups obtained per litter from a mating between wild-type (WT) females and males that were either WT, HET, or KO. Each data point represents an independent litter resulting from the following breeding pairs: 3 (WT male X WT female), 5 (HET male X WT female), and 8 (KO male X WT female, *p-value* <0.0001). B-C) Frequency of litters born days after the start of the breeding cages of females (B) or males (C) of WT, HET, and KO genotypes with WT animals of the opposite sex. D) Number of pups obtained per litter from a mating between WT females and WT males, males heterozygous for a deletion of the *Rnase10* gene *(Rnase10* -/+), or males homozygous for a deletion of the *Rnase10* gene *(Rnase10^-/-^), p-value* = 0.0035). **** *p-value* <0.0001, ** *p-value* <0.01, ns (not significant).

A previous study reported that *Rnase10* deletion in the C57BL/6 strain background, but not in the CD1 outbred strain, leads to fertility defects [43]. The loss of fertility in KO mice in the FVB background (generated in the current study) might be due solely to the deletion of the *Rnase10* gene. We, therefore, generated mice with *Rnase10* deleted in the FVB background (**Figure S1C**). *Rnase10* knockout males produced fewer pups per litter but did not show complete loss of fertility as observed in the KO mice (**Figure 2D**). These data demonstrate that the sterile phenotype of the *Rnase9-12* knockout males is not simply a consequence of the loss of the *Rnase10* gene and that additional genes from the *Rnase9-12* cluster impact fertility.

### *Rnase9-12* knockout sperm are defective in fertilizing oocytes *in vivo* but capable of fertilizing oocytes *in vitro*

Failure of KO animals to sire litters could result from defects at multiple stages during reproduction, ranging from inability to fertilize oocytes to failure of embryos to implant to early fetal failure. Therefore, we next examined the fertilization rate of embryos produced by mating WT females with WT, HET, or KO males. Superovulated females were housed with males overnight, and the following day, oocytes were collected from the oviductal ampullae of females that displayed a copulatory plug. The oocytes were cultured *in vitro* to allow the development of fertilized oocytes until the blastocyst stage. Interestingly, while ∼60% of oocytes isolated from females mated with WT and HET males reached the 2-cell stage, only 15% of oocytes derived from mating with KO males reached the 2-cell stage; the remaining cells were primarily unfertilized oocytes. Furthermore, while embryos from WT and HET males progressed well to later stages of development, with approximately 50% reaching the blastocysts stage, most fertilized KO embryos did not progress beyond the 2-cell stage, and only 3% reached the blastocysts stage (**Figure 3A**). These data showed that KO sperm were defective in fertilizing oocytes during natural mating.

**Figure 3:**
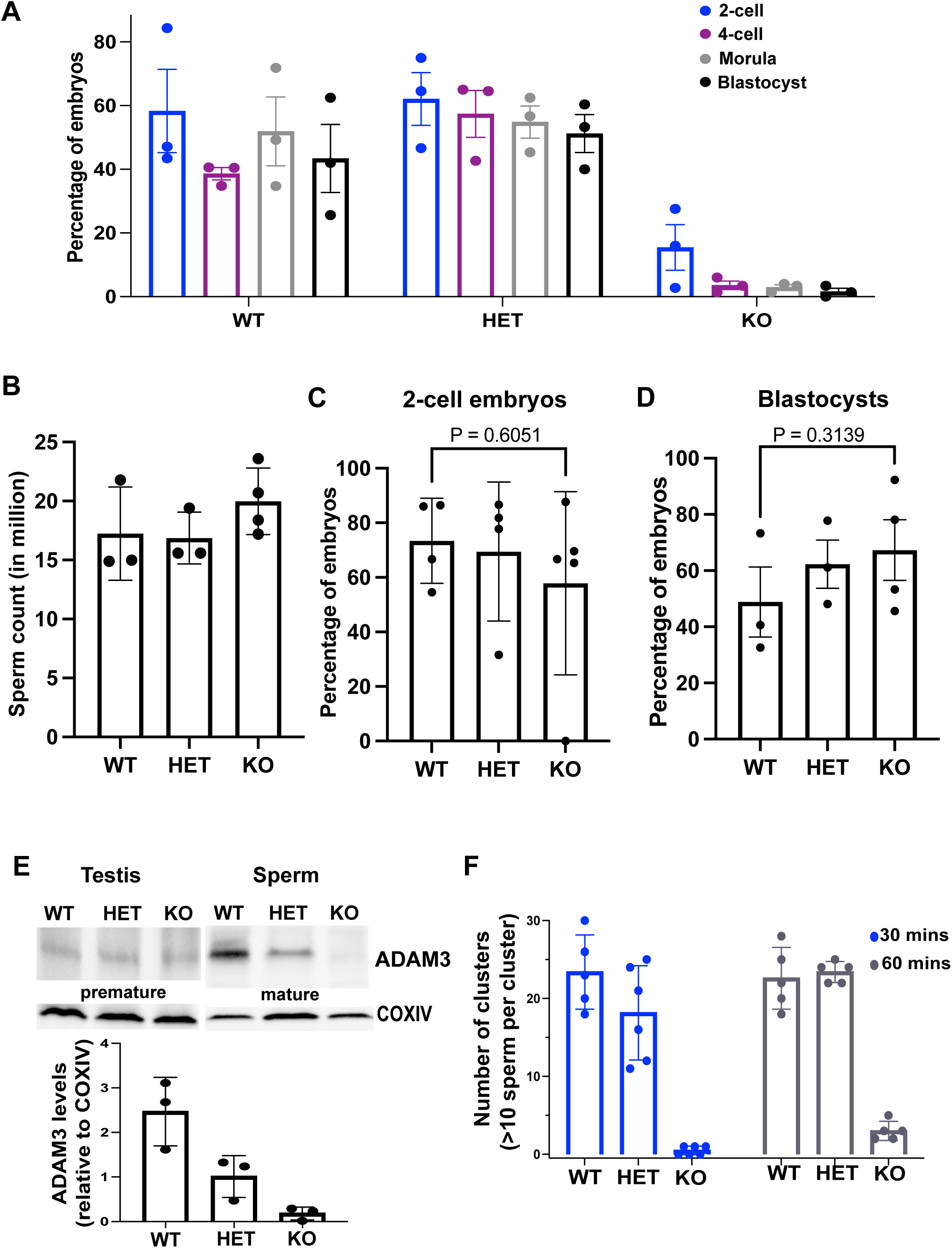
*Rnase9-12* KO sperm are capable of fertilizing oocytes *in vitro*. A) Percentage of embryos at specific preimplantation developmental stages. WT females were mated with WT, HET, or KO males overnight, and oocytes were collected from females that showed copulatory plugs the next day. The fertilized oocytes were allowed to develop *in vitro* to the blastocyst stage, and embryos at each stage were qualified as the percent of total oocytes that reached the specific developmental stage. For example, the percentage of 2-cell embryos was calculated as a percentage of oocytes that reached the 2-cell stage. The experiment was performed with 3 males per genotype and 7-9 total females per genotype. B) Total cauda epididymis sperm count in WT, HET and KO males. C-D) Oocytes from WT females were *in vitro* fertilized using sperm from WT, HET, or KO males. The plots show the percentage of oocytes that developed to the 2-cell embryo stage (C) and the percentage of 2-cell embryos that reached the blastocyst stage (D). E) Western blot analysis of mature ADAM3 protein levels in WT, HET, and KO sperm. A premature form of ADAM3 was detected in the testis (∼100 kDa), and mature ADAM3 was detected in cauda sperm (∼30 kDa). COXIV was used as a loading control. The bar graph represents the quantification of ADAM3 levels relative to COXIV in three independent biological replicates. F) Bar graph depicting the number of sperm clusters observed in the three genotypes. Sperm aggregates (10 or more sperm per cluster) in each microscopic field were calculated in three independent biological replicates. Representative microscopic images are included in supplementary Figure S2.

We next investigated the biological basis of the low fertilization rate by sperm from KO males. While the KO sperm counts were comparable to those from HET and WT males (**Figure 3B**), KO sperm displayed slight to moderate motility defects across different males (n=3) (**Supplementary Movies 1-6**). To examine the ability of KO sperm to fertilize embryos *in vitro*, we performed *in vitro* fertilization (IVF) using oocytes collected from WT females and sperm collected from WT, HET, or KO males. Interestingly, 2-cell embryos were obtained at comparable rates in all three conditions (there was a lower number of KO sperm-derived 2-cell embryos, but the difference was not statistically significant) (**Figure 3C**), demonstrating that KO sperm can fertilize oocytes *in vitro* but fail to do so efficiently during natural mating. Furthermore, we did not detect any significant difference in the percentage of blastocysts obtained from IVF using KO sperm compared to WT and HET sperm (**Figure 3D**). This implies that preimplantation development was not affected in KO sperm-derived embryos generated via IVF.

It was previously reported that *Rnase10* deletion mice have fertility defects due to failure in sperm transit through the utero-tubular junction (UTJ) [43]. Given that KO sperm fertilized oocytes *in vitro* and did not show any observable defects in sperm morphology and only a subtle motility defect, we examined whether KO sperm are defective in migrating through the UTJ. The inability of sperm to pass through the UTJ can be caused by the loss of ADAM3, a disintegrin and metallopeptidase transmembrane protein, from the surface of cauda epididymal mature sperm [48]. ADAM3 is expressed as a precursor protein in the testis and gets processed to generate a smaller mature protein during the epididymal maturation of sperm. While ADAM3 precursor protein levels in the testis were comparable between the three genotypes, KO sperm had no detectable mature ADAM3 protein (**Figure 3E**). Notably, the levels of ADAM3 were also decreased in HET mice, but the loss was not as dramatic as observed in KO sperm and was sufficient to support fertility. Moreover, KO mice showed a sperm aggregation defect; while WT and HET sperm formed clusters after release from the epididymis, KO sperm did not form clusters (**Figure 3F and Figure S2**). Sperm aggregation defects are also observed in *Adam3* [49] and *Rnase10* [43] knockout mice, suggesting that this phenotype is linked to ADAM3 loss. Together, these data imply that KO sperm are defective in transiting through the UTJ [50].

### Transcriptomic and proteomic alterations in *Rnase9-12 KO* epididymis tissues

We performed transcriptome analysis on the epididymis of KO and WT mice to investigate potential pathways affected in KO mice. As expected, *Rnase9-12* were the most significantly downregulated transcripts in KO epididymis tissues compared to WT tissues **(Figure S3A-C).** There were 28, 330, and 113 significantly differentially expressed transcripts in the KO caput, corpus, and cauda epididymis relative to WT (*padj* value <0.05, Log2Fold change >0.5) (**Table S1**), respectively. In the caput, all 28 misregulated genes were downregulated in the KO. In the corpus, roughly half (167/330) of the misregulated genes were downregulated. And in the cauda, 89 of the 113 misregulated genes were downregulated (**Figure 4A**). Notably, several downregulated genes in KO epididymides have been previously reported to regulate sperm motility and male fertility, including *Spint4*, *Spint5*, *Crisp1*, *Spinkl*, and *Kcnj16* [51–54].

**Figure 4:**
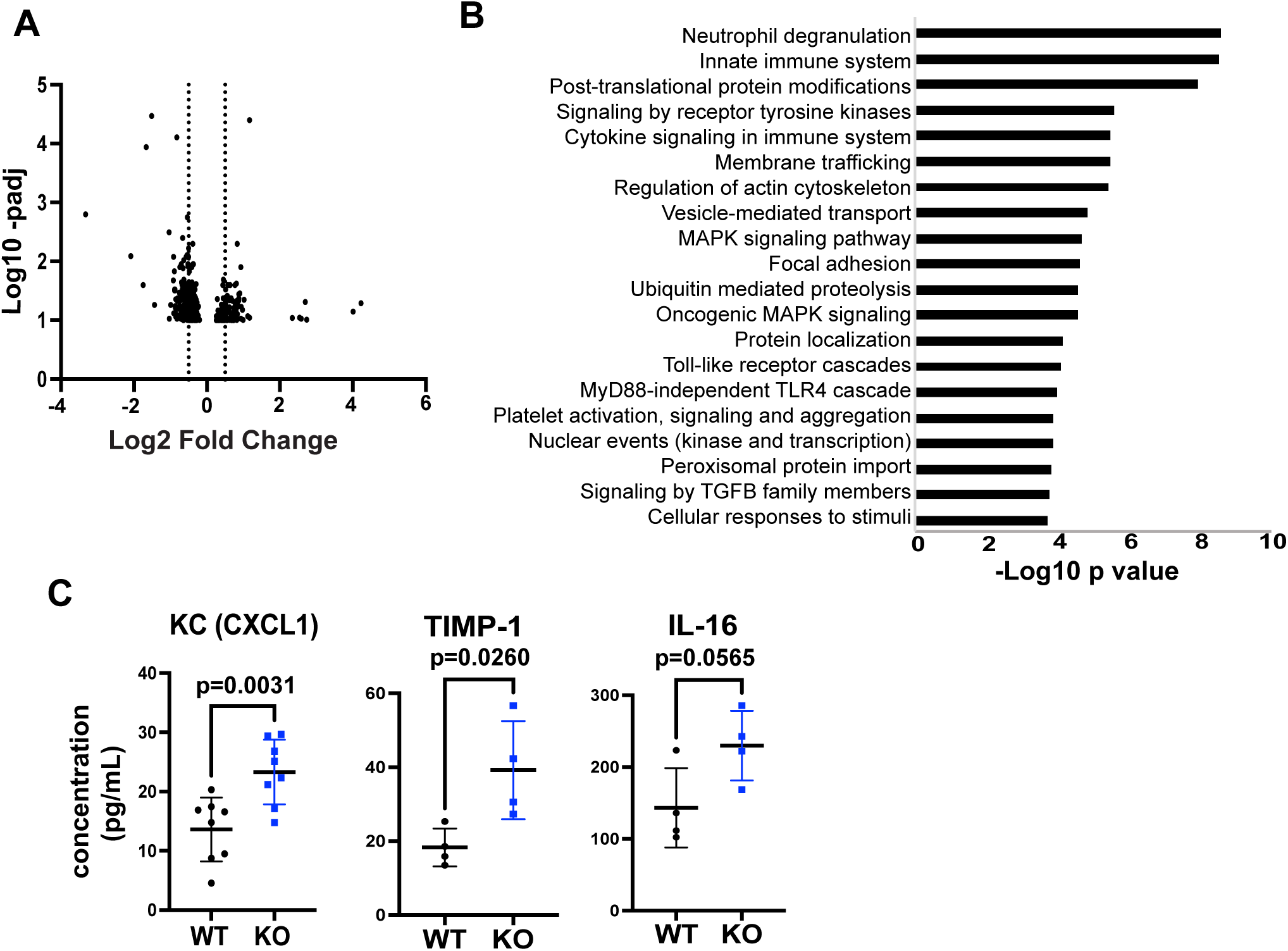
Transcriptomic and proteomic alterations in *Rnase9-12* KO epididymides. A) Volcano plot showing mRNA-seq results from DESeq2 analysis of significantly differentially expressed transcripts in the KO cauda epididymis relative to the WT (n=3 biological replicates). As *Rnase9-12* genes showed the highest and most significant changes in mRNA abundance, those four genes were removed from this plot to make other changes discernable. Volcano plots with all genes in all three regions of the epididymis are included in Figure S3. B) Gene Ontology Reactome Pathway enrichment analysis of significantly downregulated genes in cauda epididymis tissue. C) Cytokine and chemokine levels in the cauda epididymal fluid of WT and KO mice.

Gene ontology Reactome pathway analysis of significantly downregulated genes in the cauda epididymis revealed that the top pathways overrepresented were immune system related, including neutrophil degranulation, innate immune system, cytokine signaling in the immune system, and Toll-like receptor cascades (**Figure 4B**). In addition, genes involved in translation, cell adhesion, and intracellular transport were over-represented in the differentially downregulated gene set (**Figure 4B**). To determine if immune processes are altered in KO mice, we examined the levels of cytokines and chemokines in WT and KO cauda epididymides and epididymal luminal fluid. While we did not observe any significant changes in cytokine levels in the epididymis epithelial cells (data not shown), there was an increase in proinflammatory chemokine KC (CXCL1) and cytokines IL-16 and TIMP-1 in KO epididymal fluid compared to WT fluid (**Figure 4C and Figure S3D).**

Proteomic analysis using LC-MS revealed that 109, 39, and 23 proteins were significantly differentially expressed (student’s t-test, p-value<0.05) in the caput, corpus, and cauda epididymis of KO mice relative to WT, respectively (**Table S2**). Most misregulated proteins were downregulated in the KO epididymis relative to the WT epididymis tissues. Across all three segments of the epididymis, proteins that are part of the 26S proteasome complex were downregulated, including PSMB4, PSMB5, PSMB6, PSMC3, PSME1, and PSME3. Downregulation of the 26S proteasome complex has been associated with male normozoospermic infertility in humans [55] and is involved in proper sperm capacitation and acrosome reaction [56]. In addition, proteins involved in the initiation of translation and rRNA processing were downregulated, including RPL23, RPL13, RPL23a, RPL22, RPS28, RPS23, RPS21, and RPLP0. Levels of proteins involved in sperm energy generation and mitochondrial function were also reduced, such as CBR4, DLTA, PDHA2, MLYCD, IDH3A, SUCLG1, UQCRQ, UQCRSF1, COX7A2L, NDUFB8, OXSM, PGLS, ACOX1, ACOX3, L2HGDH, and PDK3. TOLLIP and STAT3, known to have anti-inflammatory functions, were downregulated in KO epididymides. Notably, antioxidants, such as GCLM and NQO1, were downregulated, suggesting that oxidative stress might result in inflammation. These data demonstrate a broad impact of *Rnase9-12* deletion on the biology of the epididymis, including a potential effect on the immune response.

### Altered levels of small RNAs in the epididymis of *Rnase9-12 KO* males

RNase A family members have been implicated in the processing/cleavage of noncoding RNAs. For example, Angiogenin cleaves tRNAs to generate tRFs in response to stress [26] and inflammation [29]. RNases 9-12 are predicted to be catalytically inactive based on their sequences lacking the RNase A family-specific catalytic residues [32]. However, whether RNases 9-12 play a role in RNA processing has not been directly tested *in vivo* except for RNase10 [57]. Cleavage of mature tRNAs and rRNAs leads to the accumulation of tRNA and rRNA fragments (tRFs and rsRNAs) that are detectable by small RNA sequencing. To examine whether tRNA or rRNA processing is affected in the KO mice, we sequenced small RNAs after pretreatment with PNK, allowing us to capture species with 3’ ends (3’P, 2’3’cP) characteristic of RNase A cleavage products [58, 59]. Small RNA sequencing reads were analyzed using the tRNA Analysis of eXpression (tRAX) analytical pipeline [60] (**Materials and Methods**).

Consistent with our previous studies [8, 9], we detected diverse classes of small RNAs in the epididymis epithelial tissue (**Figure S4A**). Interestingly, the KO cauda epididymis tissues had an overall downregulation of the percentage of tRF and rsRNA reads when compared to WT tissues (**Figure 5A**). Differential gene expression analysis revealed that the most significantly altered small RNAs in the KO cauda epididymis relative to the WT were tRFs and rsRNAs, which were mostly downregulated (**Figure 5B**). Similarly, downregulation of numerous tRFs and rsRNAs was observed in the caput and corpus epididymis, albeit fewer small RNAs showed significant changes in these tissues (**Figure S4B-C**). Examining changes in abundance of all tRFs (5’, 3’, and other) and rsRNAs, we observed an overall decrease in the KO cauda epididymis relative to the WT (**Figure 5C**). Northern blot analysis also revealed reduced levels of a 5’fragment of tRNA-Valine and rsRNAs derived from 5S rRNA in KO cauda epididymides (**Figure 5D-E**). Focusing on tRFs, we found that numerous tRFs previously described to be highly abundant in the epididymis and mature sperm were downregulated, including fragments of tRNA-Gly-GCC, tRNA-Gly-CCC, tRNA-Val-CAC, tRNA-Val-AAC, tRNA-Gln-TTG and tRNA-Glu-TTC (**Figure 5F**) [9, 61]. Moreover, we observed an overall increase in longer (>40 nts) RNA molecules for both tRF reads and all other small RNA (primarily rsRNA) reads in the KO cauda epididymides compared to WT (**Figure S4D**). Although the small RNA sequencing method used here only captures a fraction of full-length tRNAs, consistent with the hypothesis that the increased levels of longer RNAs in KO cauda epididymides are due to less tRNA cleavage, we detected subtle upregulation of full-length tRNAs in the KO cauda epididymides (**Figure 5B and S4E).** Together, these data demonstrated that KO epididymides had reduced levels of tRFs and rsRNAs, suggesting that loss of the *Rnase9-12* gene cluster causes reduced biogenesis and/or reduced stability of those two classes of RNA fragments.

**Figure 5:**
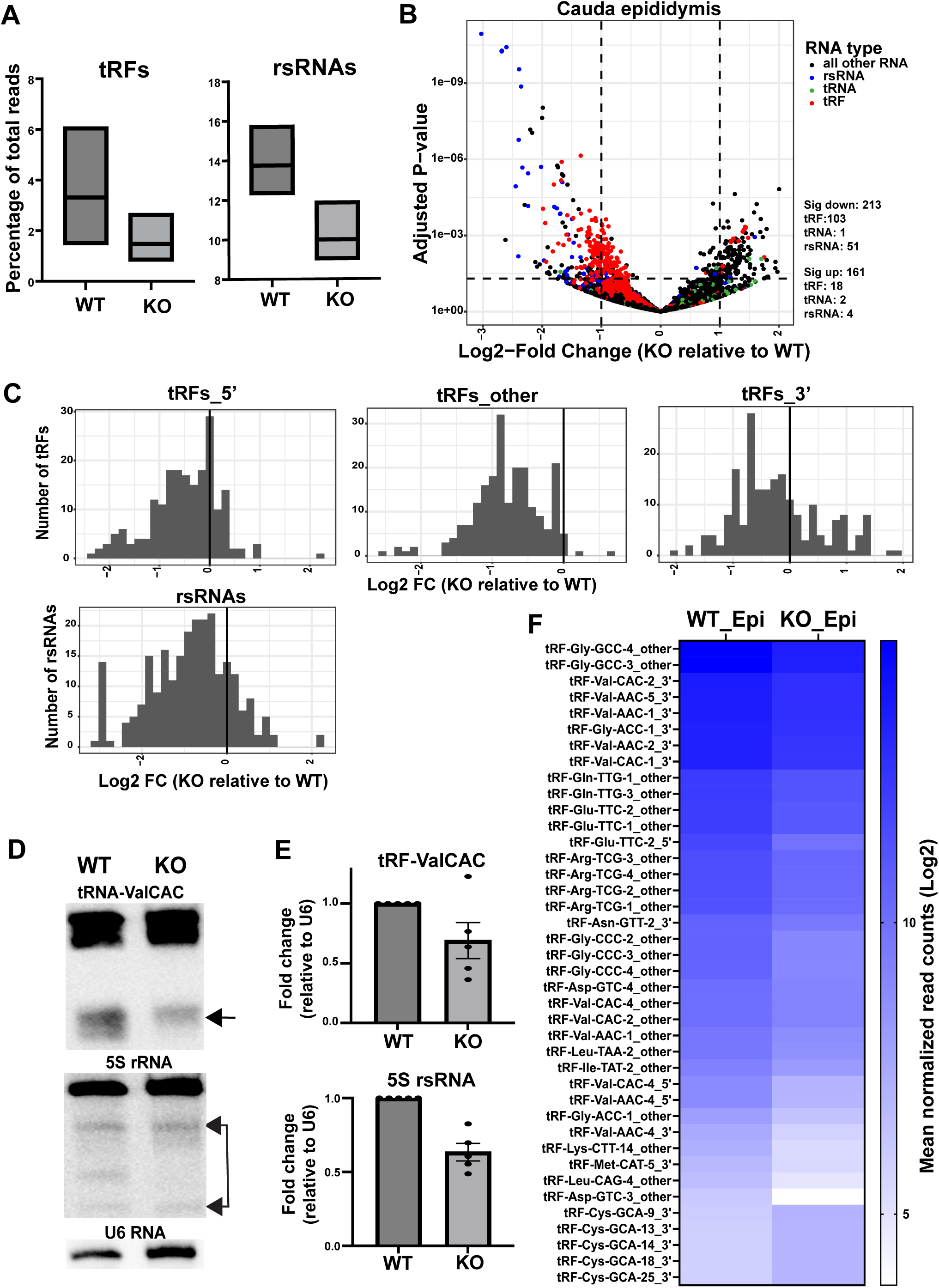
Small RNA abundance changes in the *Rnase9-12* KO epididymis. A) Percentage of tRF and rsRNA reads in the cauda epididymis tissues of WT and KO mice (n=3 biological replicates). B) Volcano plot showing differentially expressed small RNAs in the KO cauda epididymis relative to the WT cauda epididymis. tRFs are labeled red, tRNAs are green, rsRNAs are blue, and all other RNAs are black. The dashed lines show the cut-off used for calling significantly differentially expressed transcripts. The number of significantly differentially expressed transcripts (Log2Fold change >1 and padj value <0.05) as determined by DESeq2 analysis are shown in the right panel. C) Histograms showing tRF (tRF_5’, tRF_3’ and tRF_other) and rsRNA abundance changes in KO cauda epididymis relative to the WT. The X-axis shows the log2 fold change of the median of normalized reads, and the Y-axis shows the total count of tRFs or rsRNAs. D) Northern blot analysis of fragments derived from tRNA-ValCAC and 5S rRNA. Arrows indicate the fragments that were quantified. E) Quantification of tRF-ValCAC and 5S rsRNA levels relative to U6. F) Heatmap showing log2 mean of normalized read counts of top tRFs significantly differentially expressed between KO and WT cauda epididymides.

RNase A family proteins are secreted extracellular proteins. RNase1 was recently shown to regulate levels of extracellular tRFs and Y-RNA fragments [34]. Moreover, porcine orthologue of *Rnase10* was identified as the most abundant secreted protein in the proximal epididymis [62]. Therefore, we investigated if the extracellular RNA (exRNA) landscape is altered in KO mice. We isolated RNA from epididymal luminal fluid after removing sperm, extracellular vesicles, and any other cellular fractions from the fluid. Small RNA bioanalyzer analysis of exRNAs showed that caput and corpus epididymal fluid contained an almost equal proportion of 20-40nts and 70-80nts size RNA peaks. In contrast, cauda epididymal fluid had a higher abundance of 20-40nts size RNAs **(Figure S5),** suggesting increased RNA processing or higher stability of <40nt size RNAs in the distal cauda epididymal fluid. Intriguingly, small RNA-sequencing revealed that epididymal exRNAs were highly enriched in tRFs (**Figure 6A**). These findings are consistent with recent reports demonstrating that tRFs are typically more stable in cell culture media than most other RNA species [63]. Similar to epididymis epithelial cells, we observed a downregulation of the percentage of tRF reads in the KO epididymal fluid compared to WT (**Figure 6A**). Differential gene expression analysis revealed altered levels of tRFs and rsRNAs in the KO caput, corpus, and cauda epididymal fluid relative to WT, with the highest number of differentially expressed transcript in the corpus epididymal fluid (**Figure 6B and S6A-B**). The top differentially expressed tRFs in the corpus epididymal fluid were downregulated in the KO fluid across all segments (**Figure 6C**). Moreover, as observed in the epididymis epithelial cells, caput and corpus epididymal fluid displayed increased levels of full-length tRNAs (**Figure 6D and S6D**). We note that other small RNAs were also significantly altered in the KO epididymal fluid; for example, there was an upregulation of Rny3 and Rn7sk fragments in the KO epididymal fluid (**Figure 6B and S6E**).

**Figure 6:**
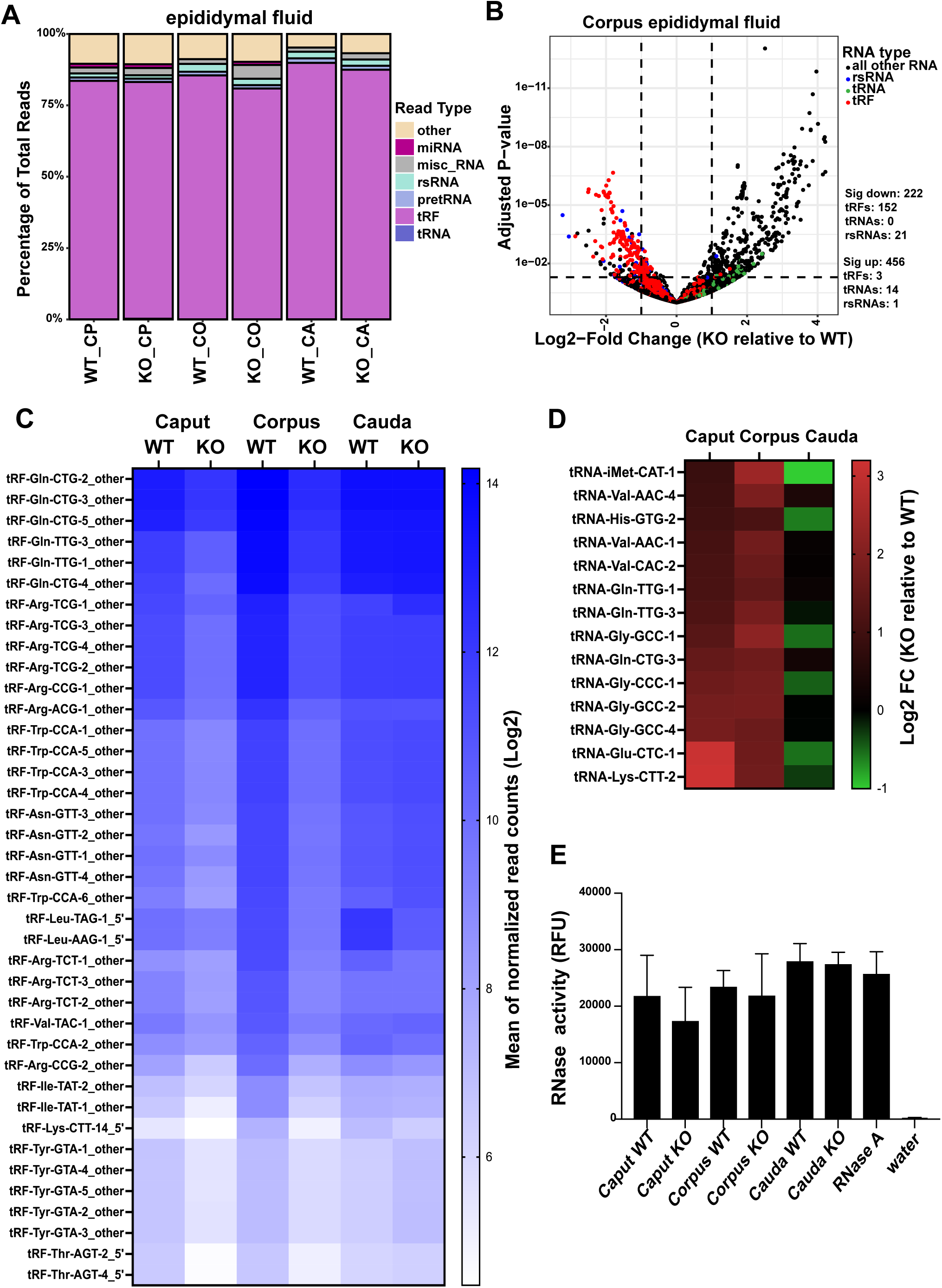
Reduced levels of tRFs in the epididymal luminal fluid of *Rnase9-12* KO mice. A) Percentage of reads of different RNA classes sequenced from the epididymal fluid (n=3 biological replicates). tRFs are fragments mapping to tRNA genes and are further classified for downstream analysis as a 5’, 3’, or “other” fragment based on the read alignment with respect to the ends of the respective tRNA (see Materials and Methods). The epididymal fluid had a high abundance of reads mapping to tRFs (> 85%). B) Volcano plot showing differentially expressed small RNAs in KO corpus epididymal fluid relative to WT. tRFs are labeled red, tRNAs are green, rsRNAs are blue, and all other RNAs are black. The dashed lines show the cut-off used for calling significantly differentially expressed transcripts. The number of significantly differentially expressed transcripts (Log2Fold change >1 and padj value <0.05) as determined by DESeq2 analysis are shown in the right panel. C) The heatmap shows the abundance (Log2 mean of normalized read counts) of tRFs in the epididymal fluid across all segments of the epididymis. Top differentially expressed tRFs in the KO corpus epididymal fluid relative to the WT are shown. Notably, most of these tRFs are also downregulated in the KO caput epididymal fluid relative to the WT. D) Heatmap showing Log2 fold change of top differentially expressed full-length tRNAs in the KO caput, corpus, and cauda epididymal fluid relative to WT. E) RNase activity measurement in the epididymal fluid collected from WT and KO mice (n=2). Purified RNase A and water were used as positive and negative controls. The bar graph represents mean ± SEM.

Together, epididymis epithelium and luminal fluid small RNA sequencing revealed that KO epididymides had reduced levels of tRFs and rsRNAs. We did not detect differential expression of any other members of the RNase A family (in the mRNA-seq dataset), implying that the reduced levels of tRFs and rsRNAs in KO epididymides were not due to a decrease in the expression of catalytically active canonical ribonucleases. Moreover, we did not detect a significant drop in the overall RNase activity of the KO epididymal fluid compared to WT (**Figure 6E**), indicating that other catalytically active RNases are present in the epididymal fluid.

### The profile of sperm small RNAs is altered in KO mice

Previous studies demonstrated that epididymis shapes the small RNA payload of mature sperm [8, 9, 13, 15–17, 64]. Therefore, we examined the small RNA profile of mature sperm isolated from the cauda epididymis. As previously reported, rsRNAs and tRFs were the most abundant classes of small RNAs sequenced from mature sperm (**Figure S7A**) [8, 11]. Consistent with changes in the epididymis epithelial cells, there was a reduction in the percentage of tRF reads in KO sperm compared to WT sperm (**Figure 7A and S7A**) and an increase in the abundance of >40nt RNAs (**Figure S7B**). Differential gene expression analysis revealed that numerous tRFs and a subset of rsRNAs were significantly downregulated in KO sperm relative to WT sperm (**Figure 7B and D**). Moreover, the abundance of most tRFs (5’, 3’, and other) decreased in KO sperm relative to WT sperm (**Figure 7C**), implying a global downregulation of tRF levels in KO sperm. Notably, other small RNA classes, such as miRNAs, did not show similar downregulation in KO sperm (**Figure 7C and S7C)**. Together, these studies showed that KO sperm have altered small RNA composition and implicate *Rnase9-12* in regulating sperm tRF levels. Importantly, these studies demonstrate the role of epididymis-expressed RNase A family members in regulating sperm small RNA profile and highlight the significance of sperm maturation in the epididymis in shaping the small RNA payload of mature sperm.

**Figure 7:**
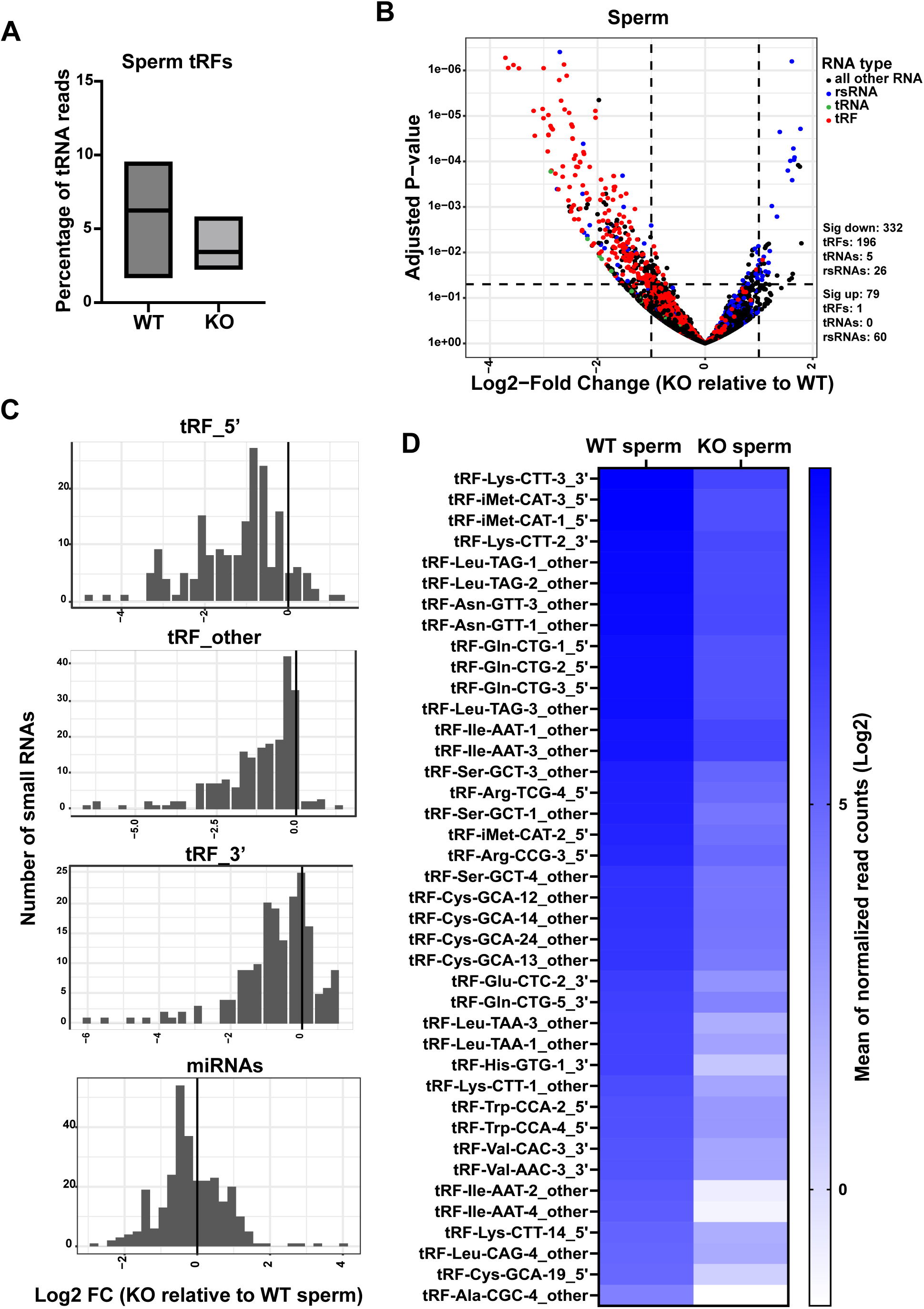
Changes in small RNA abundance in sperm from KO mice. A) Percentage of reads that mapped to tRFs in WT and KO sperm (n=4 biological replicates). B) The volcano plot shows differentially expressed small RNAs in KO sperm relative to WT. tRFs are labeled red, tRNAs are green, rsRNAs are blue, and all other RNAs are black. The dashed lines show the cut-off used for calling significantly differentially expressed transcripts. The number of significantly differentially expressed transcripts (Log2Fold change >1 and padj value <0.05) as determined by DESeq2 analysis are shown in the right panel. C) tRF (tRF_5’, tRF_3’ and tRF_other) and miRNA abundance change in KO sperm relative to WT sperm. The X-axis is the log2 fold change of the median of normalized reads, and the Y-axis shows the total number of tRFs or miRNAs. D) The heatmap shows the log2 mean of normalized read counts of top tRFs, which is significantly differentially expressed between KO and WT sperm.

## DISCUSSION

### *Rnase9-12* are required for male fertility

The epididymis is essential for male fertility, but the specific factors and mechanisms underlying this regulation remain poorly understood. Here, we identified the role of four epididymis-specific RNase A family members in regulating male fertility. Deleting a genomic locus on chromosome 14 harboring the *Rnase9-12* genes resulted in a complete loss of male fertility. As the deletion of a single gene (*Rnase10)* from this cluster resulted in subfertility, the sterile phenotype observed in KO males (*Rnase9-12* deletion) is likely due to a cumulative effect of deletion of two or more genes in this cluster. Similar phenomena have been observed for other gene families [65, 66]. For example, deletion of a cluster of beta-defensins from chromosome 8 in mice resulted in sterility [67], while deletion of a single defensin gene resulted in only subtle phenotypes [68]. RNase9-12 may function in fertility by regulating sperm transit through the UTJ [43], sperm motility [44], and/or other aspects of sperm maturation in the epididymis. Given their segmented expression along the length of the epididymis (**Figure 1**) —*Rnase10* is expressed exclusively in the caput, *Rnase9* mainly in caput and corpus, and *Rnase11* and *Rnase12* in the caput, corpus, and cauda epididymis — these genes likely work cooperatively to regulate sperm maturation in the epididymis.

The sperm membrane protein ADAM3 is thought to play a pivotal role in sperm-zona pellucida binding and sperm migration through the UTJ [48, 69]. KO sperm showed loss of ADAM3 and sperm aggregation defects. We also observed subtle motility defects in KO sperm. These results suggest that loss of ADAM3 from the KO sperm surface may result in an inability of sperm to migrate through the UTJ, leading to fertility defects. However, as *Rnase10* deletion also resulted in decreased sperm ADAM3 levels and an aggregation defect [43], modulation of additional factors in the KO epididymis likely results in the sterility phenotype observed in *Rnase9-12* KO mice. To investigate the underlying molecular changes in the KO epididymis, we performed transcriptomic and proteomic analyses of the KO tissues relative to the WT. Expression of genes involved in many biological pathways was altered in the KO epididymis, including those involved in regulating fertility.

### *Rnase9-12* regulate biogenesis and/or stability of tRFs and rsRNAs in the epididymis

RNases 9-12 are members of the vertebrate-specific RNase A superfamily and are paralogous to Angiogenin [40]. Angiogenin can cleave tRNAs to generate tRFs in response to stress and inflammation [26, 29]. It was recently reported that inflammation can induce the expression of Angiogenin in the epididymis, resulting in increased levels of specific tRFs in sperm [29]. However, as robust tRNA cleavage is observed in the epididymis even in the absence of any stressors, other RNase A family endonucleases likely contribute to tRF biogenesis under non-stress, physiological conditions. Therefore, we focused on four paralogs of Angiogenin — *Rnase9-12*— that are highly and specifically expressed in the epididymis. Here, we found that deletion of these four epididymis-specific RNase A family members led to a dramatic decrease in the levels of tRFs and rsRNAs in the epididymis and sperm, suggesting a reduced cleavage of their precursor RNA molecules (tRNAs and rRNAs) and/or decreased stability of tRFs and rsRNAs in the KO epididymis. Together, these observations underscore the roles of RNases 9-12 in regulating small RNA levels in sperm.

Numerous studies have reported that tRFs and other noncoding RNAs are abundant in bodily fluids. These extracellular RNAs can be encapsulated in extracellular vesicles (EVs) and lipoprotein particles or as free-floating RNAs in ribonucleoprotein complexes. Recent reports demonstrated the role of RNase 1 in cleaving tRNAs and Y-RNAs present in the non-vesicular extracellular environment of cultured human cells [34]. Since RNases 9-12 belong to the same family of enzymes, we investigated their role in processing exRNAs in the epididymal luminal fluid. Intriguingly, small RNA sequencing revealed that in the epididymal fluid, >85% of reads were from tRFs compared to less than 25% of tRF reads in the epididymis epithelial cells. A high abundance of tRFs in the epididymal fluid suggests that tRNA cleavage takes place in the epididymal fluid to generate tRFs, which can be taken up by the epithelial cells along the length of the epididymis [70]. Alternatively, tRFs could be generated in the epididymal epithelial cells and selectively secreted in the epididymal luminal fluid. Notably, KO mice had decreased levels of tRFs in the epididymal fluid. The decreased levels of tRFs in the KO epididymal fluid could be due to reduced cleavage of tRNAs in the KO epididymal fluid or a reduction in the levels of tRFs secreted from the epididymis epithelial cells. An important future goal is to elucidate the functional significance of extracellular tRFs in the epididymal fluid.

RNases 9-12 are predicted to lack catalytic activity due to the absence of the conserved RNase A family catalytic residues [32], raising the question of how *Rnase9-12* regulate tRF and rsRNA levels. Recently, a bacterial endonuclease was discovered that adopts the same structural folds as Angiogenin and other RNase A paralogs but has no sequence similarity with RNase A and lacks the conserved disulfide bonds and the catalytic triad of RNase family members [45]. This protein was shown to possess potent RNase activity *in vitro* and *in vivo* despite having a catalytic core distinct from that found in the canonical RNase A enzymes. These observations suggest that paralogs of Angiogenin that lack RNase A catalytic residues could still have catalytic activity. We note that we did not detect a significant drop in the overall RNase activity in the epididymal fluid upon deletion of the *Rnase9-12* genes. This data suggests that RNases 9-12 are not the major contributors of RNase activity in the epididymis, and if they have weak RNase activity (similar to Angiogenin), it was not detected by the assay. As an alternative possibility to RNase 9-12 being capable of RNA cleavage, they could stabilize tRFs and rsRNAs by directly binding to these cleaved RNAs [71]. Finally, RNase A family members can be internalized by cells via endocytosis or dynamin-independent uptake, where they can regulate cellular processes in a catalytic-activity-dependent and independent manner [72–74]. Future studies with mice expressing epitope-tagged versions of RNases 9-12 will allow direct analysis of their *in-vivo* interacting partners to further assess the role of these genes in fertility and small RNA processing. In addition, the connection between small RNA levels and sperm transit through the UTJ in the epididymis remains to be elucidated.

### *Rnase9-12* regulate sperm tRF levels

Sperm tRFs have been implicated in regulating early embryonic development and intergenerational epigenetic inheritance [18, 19]. We previously reported that a 5’ fragment of tRNA-Glycine-GCC is upregulated in sperm of mice exposed to a low-protein diet, and in preimplantation embryos, this tRF regulates transcription of a subset of genes expressed during zygotic genome activation [9]. Recently, a tRF derived from tRNA-Glutamine-TTG was reported to regulate the early cleavage of porcine preimplantation embryos [61]. Moreover, various paternal environmental conditions (such as diet, stress, and toxicant exposure) modulate offspring phenotypes, and sperm tRFs are proposed to mediate the inheritance of such paternal environmental effects. For instance, various environmental perturbations alter levels of tRFs in the sperm of mice [9, 15, 19, 29, 75–77] and humans [76, 78, 79]. Sperm tRFs are delivered to the embryo at the time of fertilization, where they can alter embryonic gene expression and offspring phenotypes. Therefore, it is critical to elucidate the biogenesis and functions of sperm tRFs. Here, we uncovered that *Rnase9-12* regulate the small RNA composition of mature mammalian sperm. These genes, thus, could be potential targets of signaling pathways that link paternal environmental conditions to offspring phenotypes. While the KO mice cannot reproduce naturally, a litter could be generated via IVF. Irrespective of the underlying mechanism of *Rnase9-12*-mediated regulation of sperm small RNAs, the KO mice provide a unique genetic model to examine the global functions of sperm tRFs in embryonic development and intergenerational transmission of paternal environmental effects, which will be a focus of future studies.

## EXPERIMENTAL PROCEDURE

### Animal husbandry and mouse lines

All animal care and use procedures were in accordance with the guidelines of the University of California Santa Cruz Institutional Animal Care and Use Committee. Mice were group-housed (maximum of 5 per cage) with a 12-hour light-dark cycle (lights off at 6 pm) and free access to food and water *ad libitum*. The pups born were weaned at 21 days of age. *Rnase9-12* and *Rnase10* knockout mouse lines were generated by Cyagen using CRISPR-Cas9 gene editing. For *Rnase9-12* KO mice, two guide RNAs were used to delete the genomic locus harboring the four genes (Figure S1A). For *Rnase10* KO mice, two guide RNAs were used to delete the 5’ half of exon 2. The knockout mice did not show any physical defects. In the *Rnase9-12* line, we occasionally noticed smaller pups and pups with short tails; however, these phenotypes did not correlate with *Rnase9-12* deletion as they were also observed in genotypically wild-type animals of this line. For genotyping, 200uL of direct PCR lysis reagent (Viagen 102-T) and 2uL of Proteinase K (from 20mg/ml stock) were added to the tail snip and incubated at 55°C overnight, followed by boiling at 90°C for 10 minutes to isolate the genomic DNA. Next, the microcentrifuge tubes were spun at 12,000g for 1 minute, and the supernatant was either directly used for PCR or DNA was further purified by ethanol precipitation. BiomixRed (Bioline BIO-25006) PCR mix was used to perform genotyping PCR. The cycle conditions used were: 94°C for 3 minutes followed by 35 cycles of 94°C for 30 seconds, 60°C for 35 seconds and 72°C for 35 seconds, and a final extension for 5 minutes at 72°C. The PCR products were then run on a 1% Agarose gel to confirm the genotype based on the length of the amplified PCR product.

### Embryo collection after natural mating

Superovulation was induced in 8 weeks old female mice by an intraperitoneal (IP) injection of 5IU Pregnant Mare’s Serum Gonadotropin (PMSG) (ProspecBio, HOR-272), followed by an IP injection of 5IU Human Chorionic Gonadotropin (hCG) (Millipore Sigma, 230734) 48 hours later. Immediately after the hCG injection, females were placed in a cage with males to allow mating. Copulatory plugs were checked 14 hours later, and oocytes were collected from the oviducts of females who displayed copulatory plugs 18 hours after the hCG injection. Oocytes with cumulus cells were transferred to a 200uL drop of KSOM media containing 3mg/ml hyaluronidase and incubated for 3-4 minutes. Oocytes separated from cumulus cells were washed and allowed to undergo further preimplantation development in KSOM media in a humidified incubator at 37°C, 5% CO2, and 5% O2 conditions.

### Sperm agglutination analysis

Cauda epididymis and Vas deferens were collected from the 8-12 weeks old WT, HET, and KO males, and sperm were released into prewarmed Human tubal fluid (HTF) media containing 0.5% BSA and incubated at 37°C and 5% CO2. Sperm count was carried out and adjusted to 2×10^5^ sperm/ml for further analysis. Sperm aggregates (with at least 10 sperm per group) were counted after incubation of sperm for 0, 30, and 60 minutes in three independent experiments using phase-contrast microscopy.

### *In vitro* fertilization (IVF)

Cauda epididymis and vas deferens sperm were collected from 8-12 weeks old WT and KO males. The tissue was placed in 1ml prewarmed HTF media containing 0.75mM Methyl-beta-cyclodextrin, and sperm was allowed to swim out by incubating the tissue at 37°C for 20-30 minutes. Sperm were counted, and 1×10^6^ sperm was then pre-incubated for 1 hour at 37°C and 5% CO2 in 200 μl drops of fertilization medium (KSOM + 1mM Glutathione) covered with sterile mineral oil. Superovulation was induced in female mice using PMSG and hCG as described above. Cumulus-oocyte complexes were collected from the oviducts of females at 13–15 h after hCG injection. These were then transferred to the 200uL fertilization drop. The oocytes were then co-cultured with sperm at 37°C under 5% CO2 for 4 hours to allow fertilization. Fertilized zygotes were washed to get rid of excess sperm and cumulus cells and cultured to later stages of development at 37°C under 5% CO2 and 5% O2.

### Sperm motility assay

10uL of sperm solution (in HTF) was mounted on a hemocytometer slide and viewed under a phase contrast microscope (Zeiss) at 10X and 20X magnification. Sperm motility videos were recorded using Zeiss Zen Pro software. At least 3 different fields were recorded per biological replicate, and 5 independent observers did blinded analysis to determine any differences in sperm motility. KO sperm showed subtle motility defects when compared to WT sperm, with the severity of the defect being variable across replicates.

### Tissue collection

Epididymis tissues and cauda sperm were collected as described previously [8]. For epididymal fluid collection, epididymides were dissected and placed in prewarmed Whitten’s media for 30 minutes at 37°C to allow the release of epididymal luminal contents in the medium. After 30 minutes of incubation, the media containing epididymal luminal contents was transferred to a 1.5 ml tube and incubator for 15 minutes at 37°C to allow tissue pieces and any cellular debris to settle. The top fraction (containing sperm and luminal fluid) was transferred to a new tube and spun for 2 minutes at 8,000 rcf to pellet sperm. The supernatant from this spin was spun at 10,000 rcf for 30 minutes at 4°C to pellet the remaining sperm and cell debris. The supernatant was then collected and spun at 100,000 g for 2 hours at 4°C to pellet extracellular vesicles. The supernatant from the last spin is the epididymal fluid free of sperm, extracellular vesicles, and any cellular debris. It was used for small RNA sequencing to examine epididymal fluid RNA composition.

### Protein extraction and immunoblotting

Total protein was extracted from caput epididymis, testis, and sperm by homogenizing the tissue/cells in ice-cold RIPA Lysis buffer (G Biosciences) with 1X Protease inhibitor cocktail and 1X Phosphatase inhibitor cocktail. Sperm was lysed by sonication at 60% power setting for ∼5 seconds, followed by incubation on ice for 10-15 minutes. Protein was quantified using BCA assay (Pierce^TM^ BCA Protein Assay Kit). 30μg total protein from tissue samples with 4X Laemmli buffer was heat denatured at 99°C for 5 minutes and ran on resolving SDS PAGE gel (Biorad 4-20% precast gel #4561096). Sperm protein samples were prepared using a low volume of lysis buffer (∼100), and ∼30 uL was run on the gel. Proteins were transferred to the PVDF membrane and incubated with a blocking solution (5% skimmed non-fat milk in 1X TBST) for 1 hour at room temperature with slow shaking. The membrane was briefly washed with 1X TBST and incubated overnight with the primary antibody dissolved in 1X TBST at 4°C on slow shaking (ADAM3 from Santa Cruz Biotechnology (sc365288) and COXIV from Cell Signaling Technology (4844s)). The next day, the membrane was washed thrice with 1X TBST (5 minutes each at RT under rapid shaking conditions) and incubated for an hour with the HRP-labelled secondary antibody in 1X TBST on slow shaking. The membrane was then developed using BioRad Clarity Western ECL substrate and imaged in a chemi-doc (Biorad). The protein bands obtained were quantified using ImageJ.

### mRNA-sequencing and data analysis

Epididymis segments were collected as described previously [8]. An Illumina TruSeq Stranded mRNA library preparation kit was used to generate mRNA-seq libraries. The libraries were sequenced on HiSeq, and differential gene expression analysis was performed using the DESeq2 analytical tool [80].

### Quantitative real-time PCR

RNA was extracted using Tri Reagent (Sigma T9424-200ML), followed by chloroform-assisted phase separation and precipitation of RNA from the aqueous phase using isopropanol. cDNA was synthesized using Superscript III RT (Thermofisher 18080093) and random hexamers (Thermofisher SO142). qPCR reactions were performed using KAPA SYBR FAST qPCR mix (Roche KK4601) and custom-designed primers for *Rnase9, Rnase10, Rnase11*, *Rnase12, and Gapdh*.

### Northern blot analysis

Northern Blotting was performed by following the previously described method using non-radioactive probes [81] in conjunction with imaging the blots using SuperSignal™ West Femto Maximum Sensitivity Substrate (Thermofisher 34094). Briefly, RNA samples were combined with 2X Gel Loading Buffer II (Thermofisher AM8546G) and denatured at 95°C for 5 minutes, followed by at least 2 minutes at 4°C or on ice. RNA samples were run on 15% acrylamide 7M urea TBE gel, and the gel was then stained with 1X SYBR gold for 10 minutes and imaged on the Bio-Rad Gel Doc XR+ machine using the corresponding Image Lab Software. The RNA was transferred from the gel to a Nylon membrane (Sigma-Aldrich SIAL-11209299001) at 4°C in a cold room using transfer stacks (from Trans-Blot Turbo RTA kit, Bio-Rad, 1704272). Next, RNA was cross-linked by incubating at 60°C for 1-2 hours in a cross-linking solution. After rinsing the membrane, it was pre-hybridized with 15 ml of prewarmed Ultrahyb buffer (Thermofisher AM8670) at 37°C for 30 minutes. The specific LNA probe was then denatured at 95°C for 1 minute and added to the hybridization buffer. The membrane was hybridized overnight on rotation. Next, the membrane was washed twice at 37°C for 15 minutes with 2X SSC with 0.1% (wt/vol) SDS, followed by two washes at 37°C for 5 minutes with 0.1X SSC with 0.1% (wt/vol) SDS. The membrane was next washed for 10 minutes with 1x SSC at 37°C. The membrane was next treated using a Chemiluminescent Nucleic Acid Detection Module Kit (Thermofisher catalog #89880, 24 ml) and imaged with the SuperSignal™ West Femto Maximum Sensitivity Substrate (Thermo Fisher Cat# 34096) kit and the Bio-Rad ChemiDocTM MP Imaging System. The same membrane was used to probe other target RNAs after stripping and re-hybridizing in the same manner as described above.

### Proteomic analysis

Caput, corpus, and cauda epididymis were harvested from mice and stored at −70°C after flash freezing in liquid nitrogen. The tissues were taken out of −70°C and immersed in lysis buffer (1% SDS, 50 mM Tris pH8.1, 10 mM EDTA pH8, protease inhibitor cocktail) and ground with a drill and disposable tips. Tissues were heated in lysis buffer for>2 minutes at 95°C to soften the tissue and assist grinding if needed. The protein concentration of lysates was determined using the Pierce™ BCA Protein Assay Kit (ThermoFisher #23225). 10 ug of protein from each sample was run on an SDS PAGE gel made using the TGX™ FastCast™ Acrylamide Starter Kit, 12% (Bio-Rad #1610174) and using a 1X running buffer made from 10X Tris/Glycine/SDS (Bio-Rad #1610772). Gels were run until the dye front had moved a few centimeters, and then the gels were stained with Bio-Safe™ Coomassie Stain (Bio-Rad #1610786). Stained proteins were cut out of the gel and shipped to the UC Berkeley Proteomics Core for mass spectrometric analysis.

### RNase activity assay

Cauda epididymis tissues were dissected from males (10-12 weeks old) and placed in prewarmed Whitten’s media (100 mM NaCl, 4.7 mM KCl, 1.2mM KH2PO4, 1.2 mM MgSO4, 5.5 mM Glucose, 1 mM Pyruvic acid, 4.8 mM Lactic acid (hemicalcium) and HEPES 20 mM) at 37°C. A couple of small incisions were made in the cauda epididymis tissues to allow sperm and epididymal fluid to be released from the tissue into the media. Media containing epididymal fluid and sperm was used for protein purification, and total protein content was quantified using the Qubit™ Protein Assay (Thermofisher Q33211). 9 μg of protein was used in each reaction of the RNaseAlert™ Lab Test Kit (Thermofisher AM1964) to examine RNase activity. Reactions were performed as per the manufacturer’s instructions, and the fluorescence signal (indicative of RNase activity) was measured using the Qubit 4 Fluorometer.

### Small RNA sequencing and data analysis

Small RNA-seq libraries were generated using ARM-Seq [82] or OTTR-seq [59]. In both cases, the total RNA was pretreated with PNK to remove 3’ phosphates. For cauda sperm and epididymal fluid samples, total RNA from 3 mice was pooled to generate one biological replicate, and 1ug total RNA from the epididymis tissue was used for epididymis libraries. Total RNA was treated with 0.5uL of T4 PNK (NEB M0201L) at 37°C for 30 minutes in a 20uL reaction volume. PNK was heat-inactivated using 0.5uL of 0.5M EDTA at 65°C for 15 minutes, followed by treatment with 3.5uL of 100mM borax or sodium tetraborate decahydrate for 30 minutes at 45°C. Next, the total volume of the reaction was raised to 400uL using nuclease-free water for phenol-chloroform-isoamyl alcohol-mediated RNA cleanup.

Small RNA sequencing libraries using OTTR-seq were generated as described by Upton *et al*. [59] and by the manufacturers of the sequencing library preparation kit (Karnateq R2201001S) with a few modifications. Briefly, 40ng PNK-treated RNA was treated with ddRTP and target primed reverse transcriptase for 2 hours, followed by heat inactivation at 65°C for 5 minutes. The reaction mixture was next incubated with phosphatase at 37°C for 15 minutes, and RNA was reverse transcribed using BoMoc RT (N-terminally truncated *B. mori* R2 Reverse Transcriptase). The cDNA was purified using the MiniElute Reaction Cleanup kit and run on a precast 10% Urea PAGE gel (BioRad #4566033) for 45 minutes at 200V. cDNA corresponding to small RNAs, including miRNAs and full-length tRNAs, was cut from the gel using Cy5 imaging, and DNA gel extraction was performed. The extracted cDNA was used for PCR reaction for 12 cycles using Q5 Polymerase and NEB multiplex indexing primers (NEB #E7600S). The PCR product was run on a 6% PAGE gel, and bands corresponding to the expected range of miRNAs to full-length tRNAs (150bp to 250bp) were cut out. The eluted PCR product was quantified using a Qubit 1X DNA High Sensitivity kit (Q33230) and analyzed on a Bioanalyzer using a DNA high sensitivity kit (#5067-4626). The final libraries were sequenced either on an Illumina HiSeq or NextSeq instrument.

For sequencing data analysis, sequencing reads were trimmed with cutadapt with adapter “-a GATCGGAAGAGCACACGTCT” and further trimming the final TRPT base with “cutadapt -u −1” and removal of the UMI with “cutadapt -u 7.” Sequencing analysis was done using the tRAX analytical analysis tool [60] with default parameters except for a minimum non-tRNA read size of 16 and using the Ensembl gene set [83], the mm10 piRNA gene set from piPipes [84], and ribosomal RNA repeats from UCSC repeatmasker [85] as the gene set. In tRAX, reads were mapped to the mouse mm10 genome combined with tRNA sequences taken from gtRNAdb [86]. To create a sequence database for mature tRNA sequences, introns were removed, CCA tails were added, and “G” base was added to the start of histidine tRNAs. tRNA reads were defined as any reads whose best mapping includes a mature tRNA sequence. The tRAX pipeline uses bowtie2 with options “-k 100 --very-sensitive --ignore-quals --np 5 –very-sensitive,” extracts all best mappings from those results, and categorizes all tRNA mappings to acceptor type-specific, decoder type-specific, or unique tRNA transcript-specific, and only reads specific to acceptor type and anticodon were used for corresponding tRNA counts.

Reads that mapped to mature tRNAs were further classified into four fragment types based on the read alignment and the ends of the tRNAs. Reads where the 5’ end lies within 3 nts and 3’ end lies with 5 nts of the respective ends on the mature tRNAs are categorized as “whole” full-length tRNAs (referred to as “tRNAs”) and reads that overlap or closely align to either the 5’ ends or 3’ end of the tRNAs are classified as “tRF_5’” and “tRF_3’”, respectively. Mature tRNA reads that do not display these features are classified as “tRF_other”. tRFs_5’, tRFs_3’, and tRFs_other are counted together as “tRFs”. We sequence relatively low levels of full-length tRNAs as the small RNA sequencing method favors cloning and sequencing of shorter RNAs (<40 nts). The total percentage of reads for full-length tRNAs was less than 0.5% across all samples and, therefore, not visible on the plots showing the percentage of reads of different small RNA classes. For rRNAs, as all-best read mappings to the UCSC repeatmasker track and Ensembl rRNAs were used, reads that can map to multiple rRNA repeats as well as the original rRNA are included as rRNA repeats. Since small RNA sequencing cannot detect RNAs longer than 100nts, all reads mapping to rRNAs and rRNA-repeats are shorter rRNA fragments and are collectively referred to as rRNA-derived small RNAs (“rsRNAs”). Reads that mapped to the genome but did not map to any annotated feature in either the Ensembl genes, piRNA gene sets, or rRNA repeatmasker were classified as “Other” read types. Adjusted p-values and log2-fold change were calculated using DESeq2 [80] with default parameters as a component of the tRAX pipeline, and plots were generated with ggplot2 and Prism.

## ACKNOWLEDGMENTS

We thank Henry Moore and Todd Lowe for help with the initial OTTR-seq analysis of the epididymal fluid small RNAs. We thank the Lowe lab, Lucas Ferguson, and Kathy Collins for help with OTTR-seq optimization. We thank all the members of the Sharma lab for their helpful discussions on this work. US is supported by NIH grant 1DP2AG066622-01 and the Searle Scholars Program, and AG is supported by a postdoctoral fellowship from the California Institute of Regenerative Medicine (CIRM) award EDUC-12759 awarded to the Institute of Biology of Stem Cells (IBSC) at UCSC. The proteomic analysis was performed at the Vincent J. Proteomics/Mass Spectrometry Laboratory at UC Berkeley, partly supported by NIH S10 instrumentation grant S10RR025622.

## AUTHOR CONTRIBUTIONS

Conceptualization US, JFS, AG; experimental design and methodology US, JFS, AG, GK, and EEL; data analysis and visualization US, SK and ADH; funding acquisition, supervision and writing US.

## DECLARATION OF INTERESTS

The authors declare no competing interests.

## Notes

### Competing Interest Statement

The authors have declared no competing interest.

