## Supplementary figures for "Epididymis-specific RNase A family genes regulate fertility and small RNA processing"

### Supplementary Figure S1

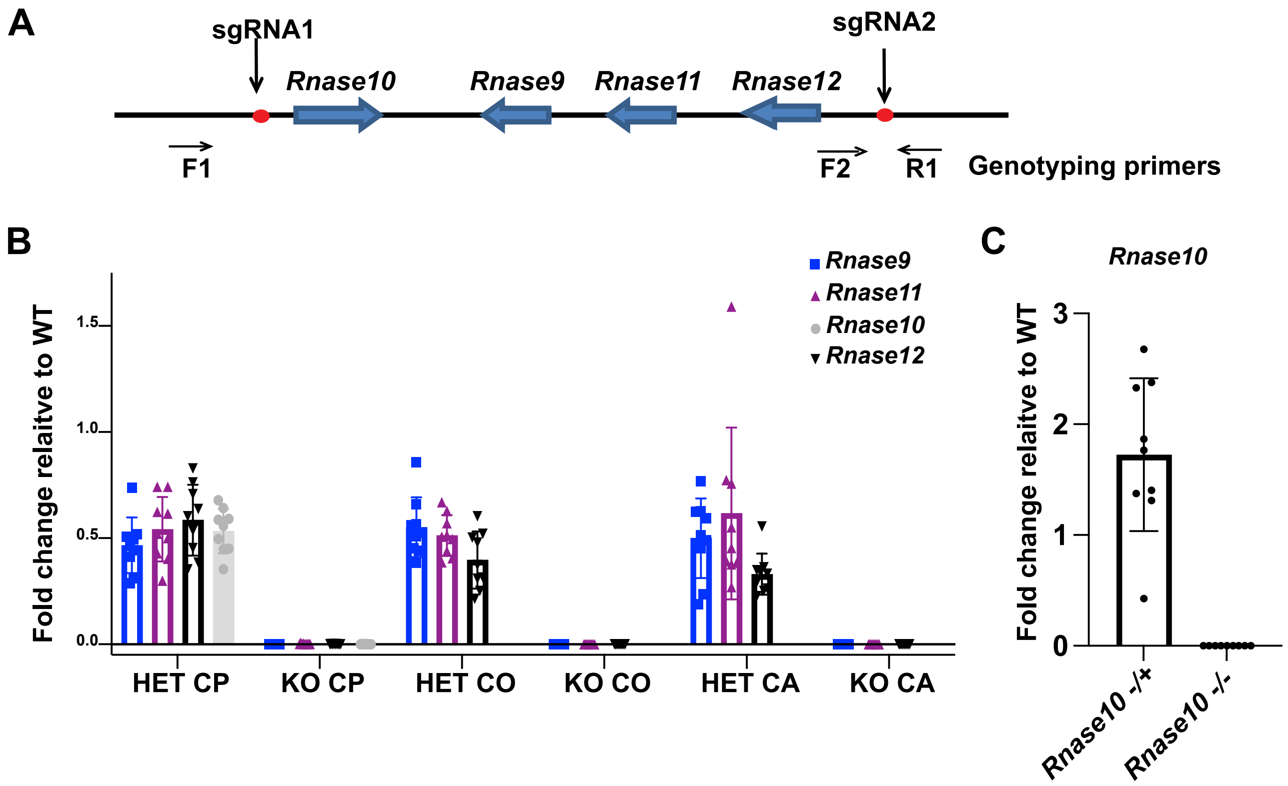

### Supplementary Figure S2

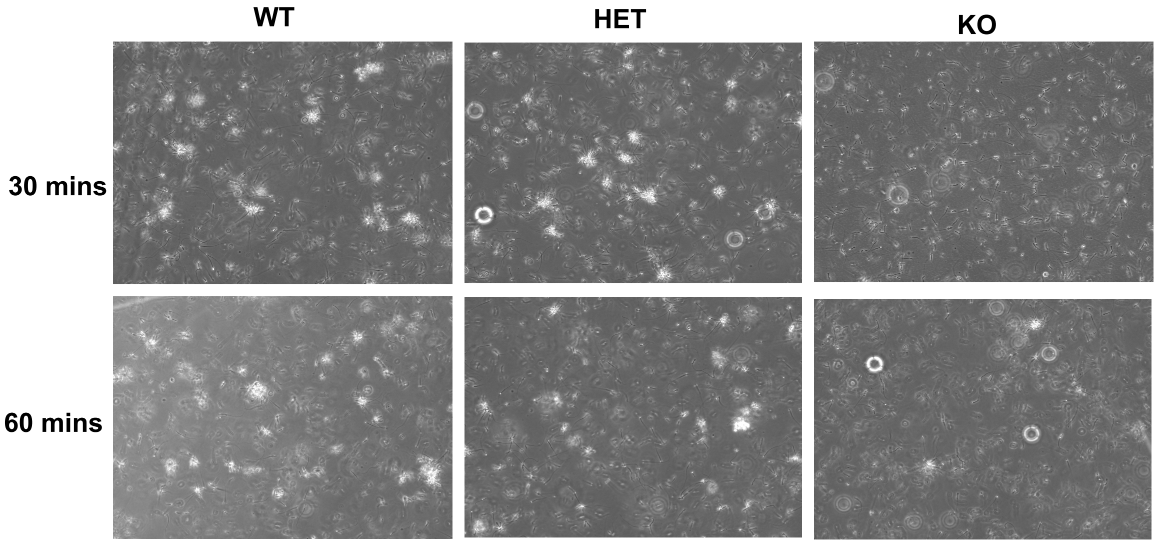

### Supplementary Figure S3

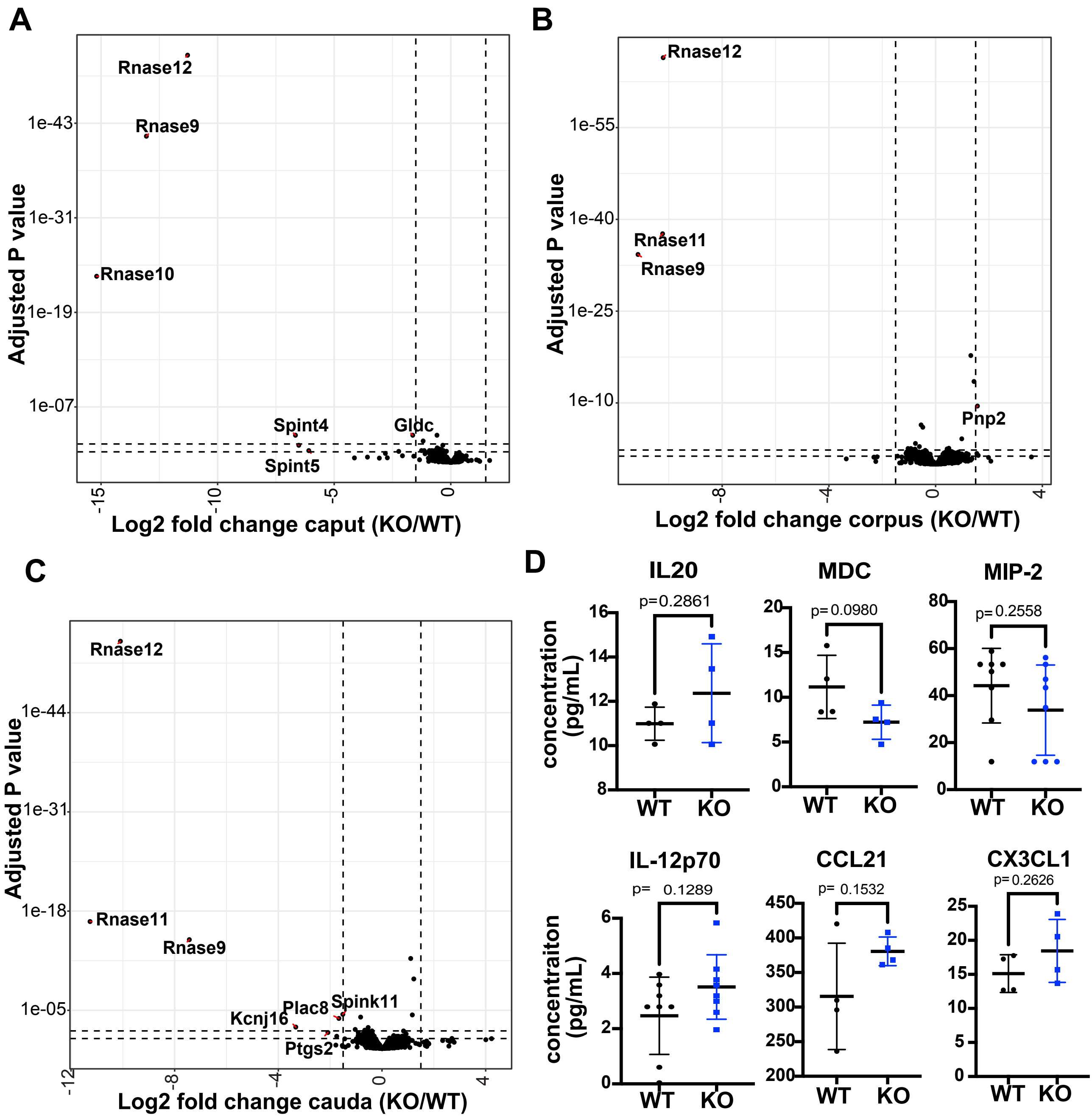

### Supplementary Figure S4

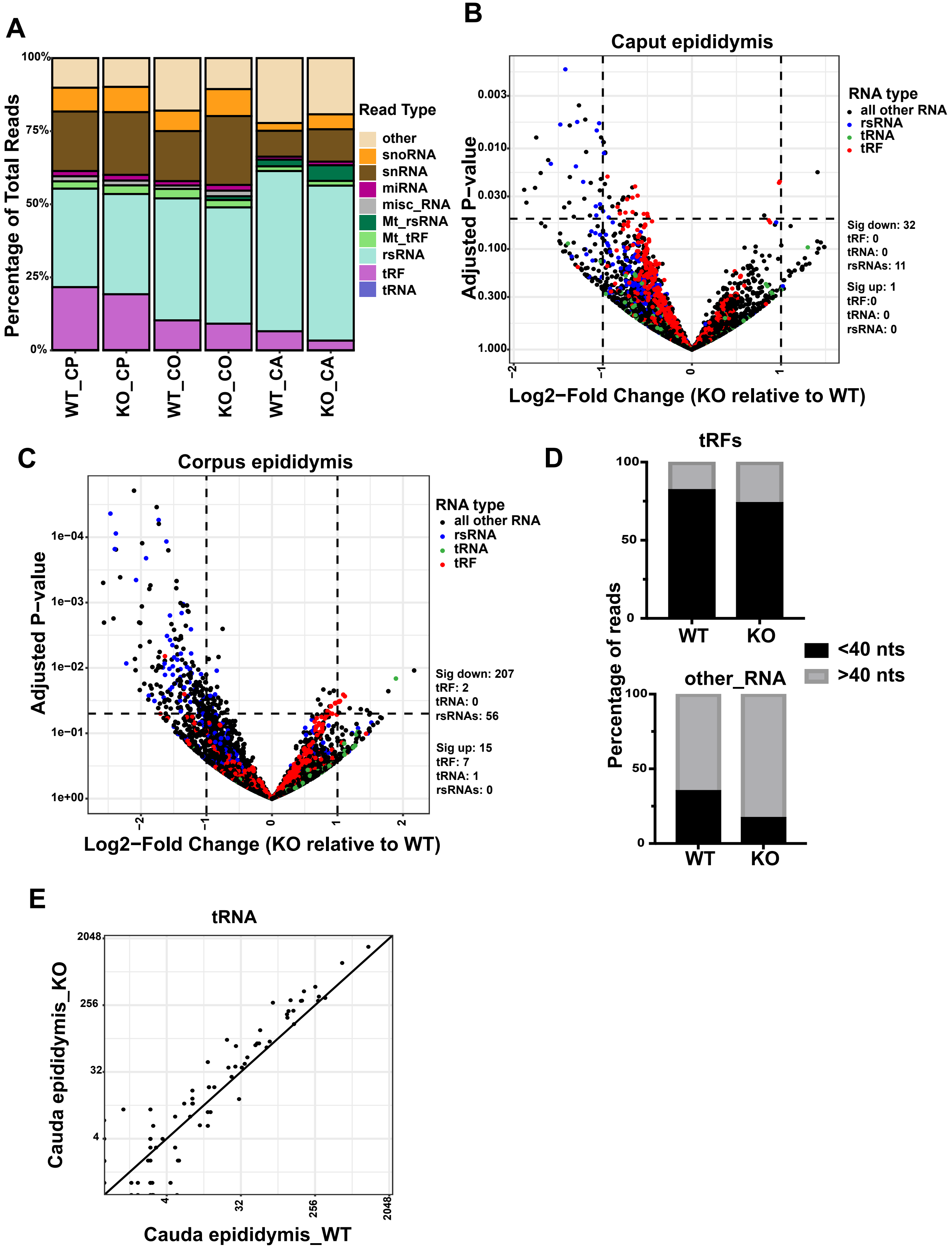

### Supplementary Figure S5

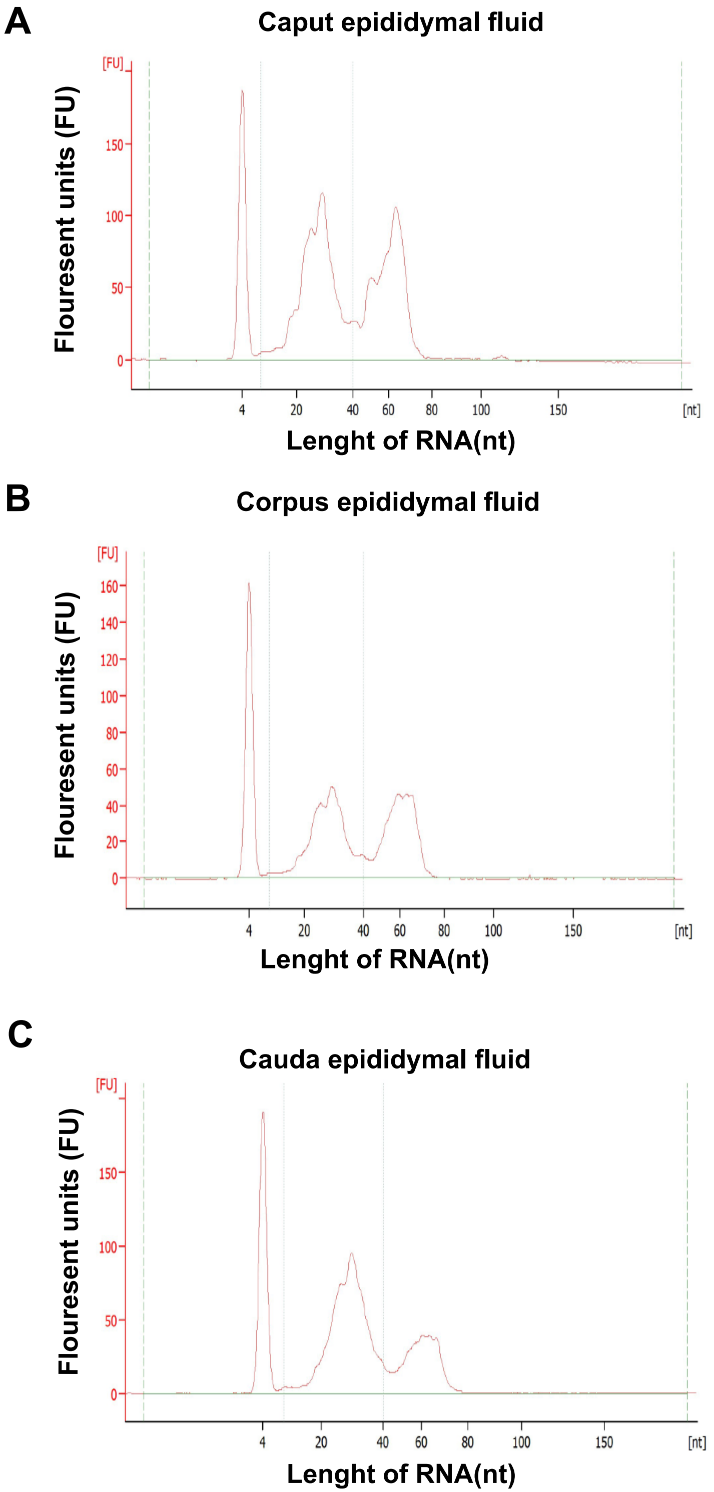

### Supplementary Figure S6

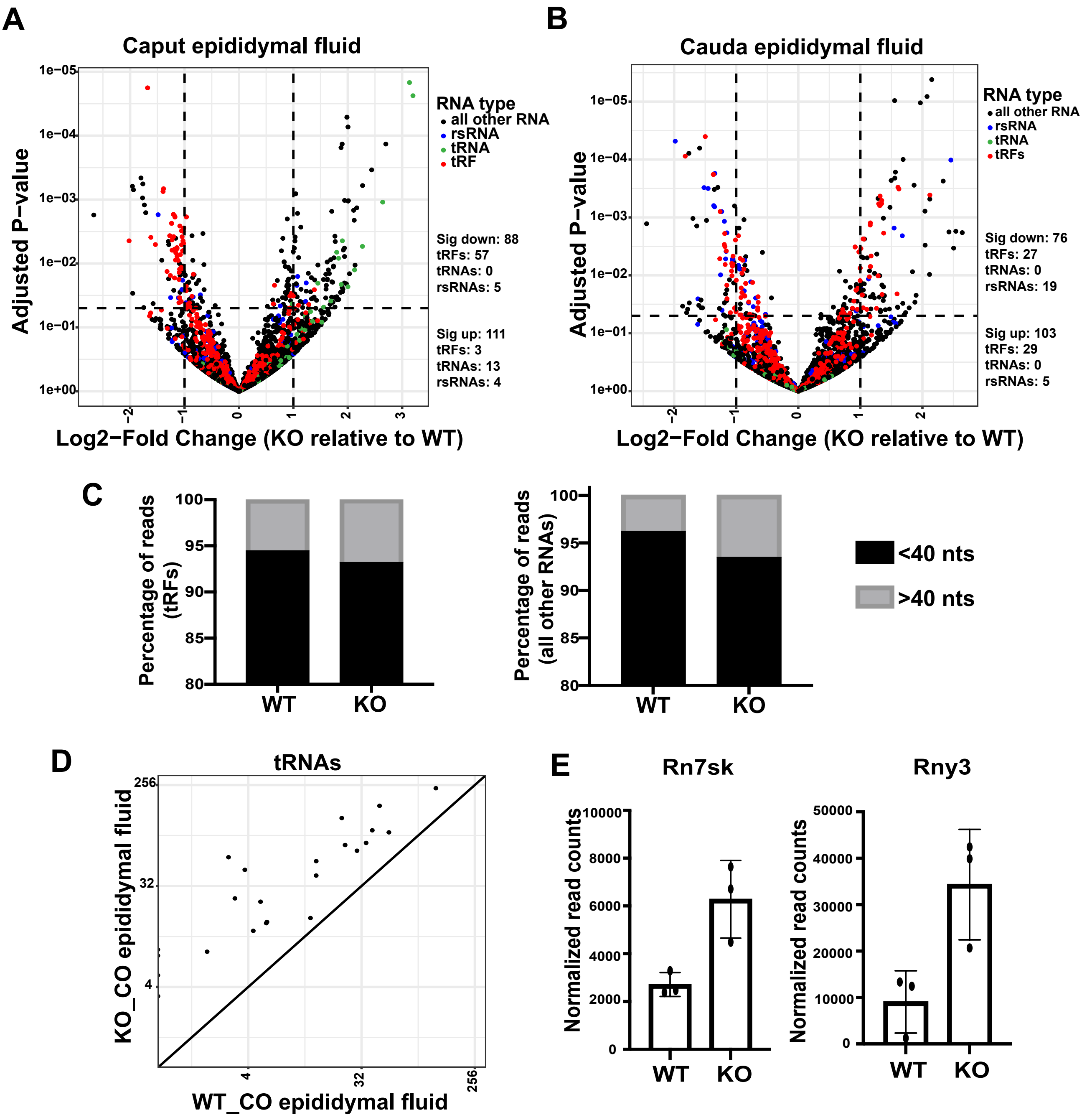

### Supplementary Figure 7

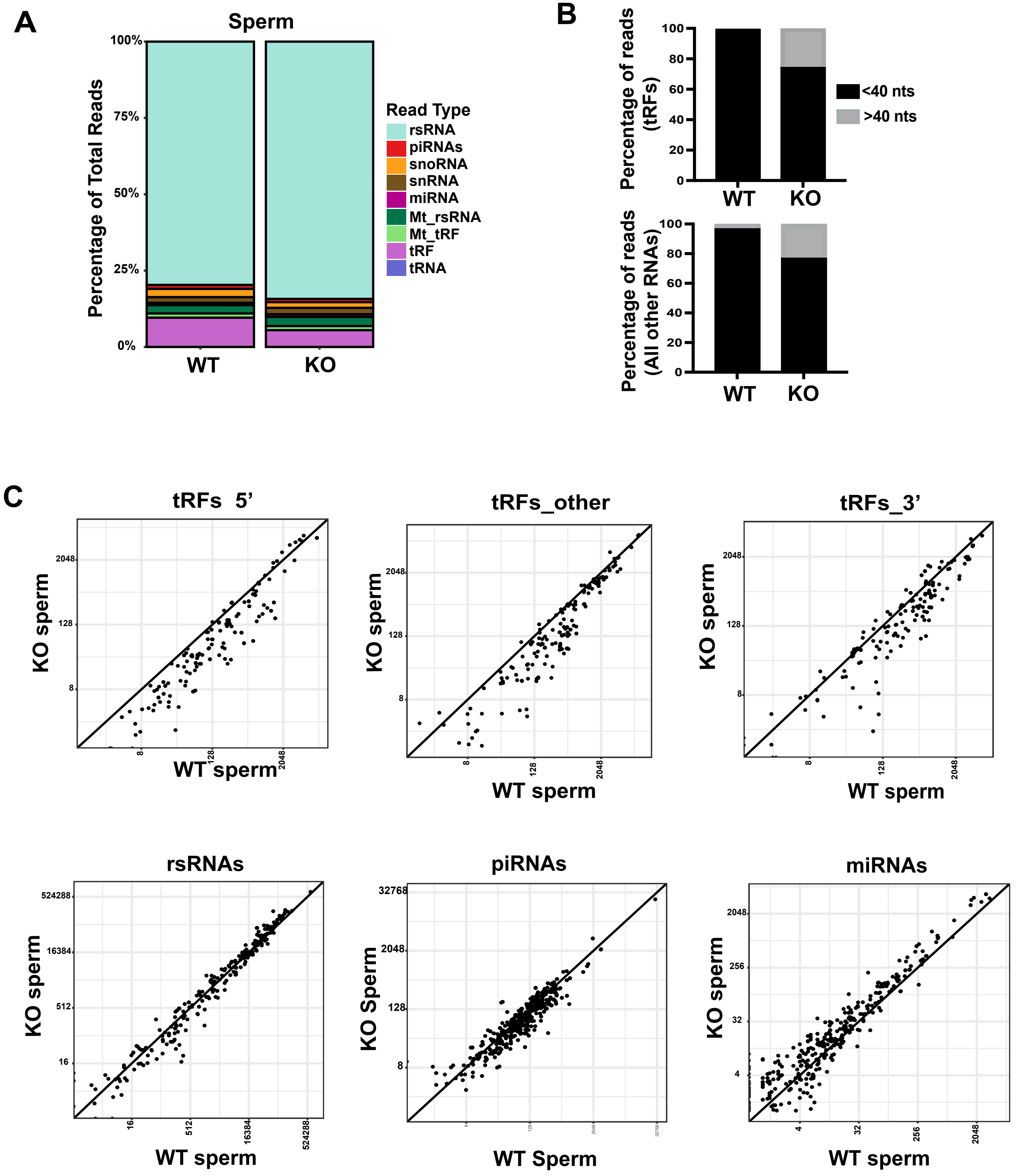
